# So close yet so far apart: Distinct flanking sequence recognition by DNMT3A and DNMT3B

**DOI:** 10.1101/2024.02.07.579311

**Authors:** Ayşe Berçin Barlas, Ezgi Karaca

## Abstract

DNMT3A and DNMT3B are closely related DNA methyltransferases that catalyze de novo CpG methylation yet exhibit distinct preferences for flanking sequences. Despite sharing 91% sequence similarity within their catalytic domains, these paralogs show non-overlapping genomic targeting and divergent biological roles. To uncover the mechanistic basis of this specificity, we performed 16µs of all-atom molecular dynamics simulations on DNMT3A and DNMT3B complexes bound to CpG substrates with varied +2 flanking bases (i.e., CGX). To resolve their base- and shape-readout mechanisms at atomistic detail, we introduce a Comparative Dynamics Analysis (CDA) framework. Our CDA analysis reveals that DNMT3A relies on a rigid, sequence-specific hydrogen bonding network and shape-constrained electrostatic anchoring, whereas DNMT3B employs a more flexible and distributed interface, allowing broader substrate tolerance. This represents the first systematic analysis of shape readout in DNMT3 enzymes and demonstrates how flanking sequence specificity is dynamically encoded by two nearly identical proteins. Our findings not only clarify how closely related DNA-modifying enzymes diverge in recognition strategies, but also lay the foundation for future efforts to engineer paralog-specific protein–DNA interactions.

## INTRODUCTION

DNA methylation is a key epigenetic modification that regulates a wide range of cellular processes, including embryonic development, gene expression, chromatin remodeling, cellular reprogramming, aging, and X chromosome inactivation ^1–4^. Unlike irreversible genetic mutations, DNA methylation is reversible, making it an attractive target for therapeutic research ^1–3^. In humans, three catalytically active DNA methyltransferases (DNMTs) are responsible for methylating cytosines, primarily at CpG dinucleotides: DNMT1, DNMT3A, and DNMT3B ^5,6^. DNMT3A and DNMT3B catalyze *de novo* methylation by targeting previously unmethylated DNA, while DNMT1 maintains existing methylation patterns during DNA replication ^7,8^. Among the *de novo* DNMT3s, DNMT3A functions in early development and adult homeostasis, whereas DNMT3B plays a complementary role, with a particular focus on centromeric and pericentromeric regions.

The paralogous enzymes DNMT3A and DNMT3B share a conserved domain architecture comprising a PWWP domain (which binds DNA and histone tails), an ADD domain (involved in protein–protein interactions), and a methyltransferase (MTase) domain that catalyzes the transfer of a methyl group from S-adenosyl-L-methionine (SAM) to the 5′ carbon of the target cytosine (Figure 1A) ^6,9–13^. The MTase domains of DNMT3A and DNMT3B are highly conserved, sharing 91.2% sequence similarity across 283 aligned residues^10^ (Figure 2A). For simplicity, from hereafter, we refer to the DNMT3A-MTase and DNMT3B-MTase domains as DNMT3A and DNMT3B throughout the manuscript.

**Figure 1.**
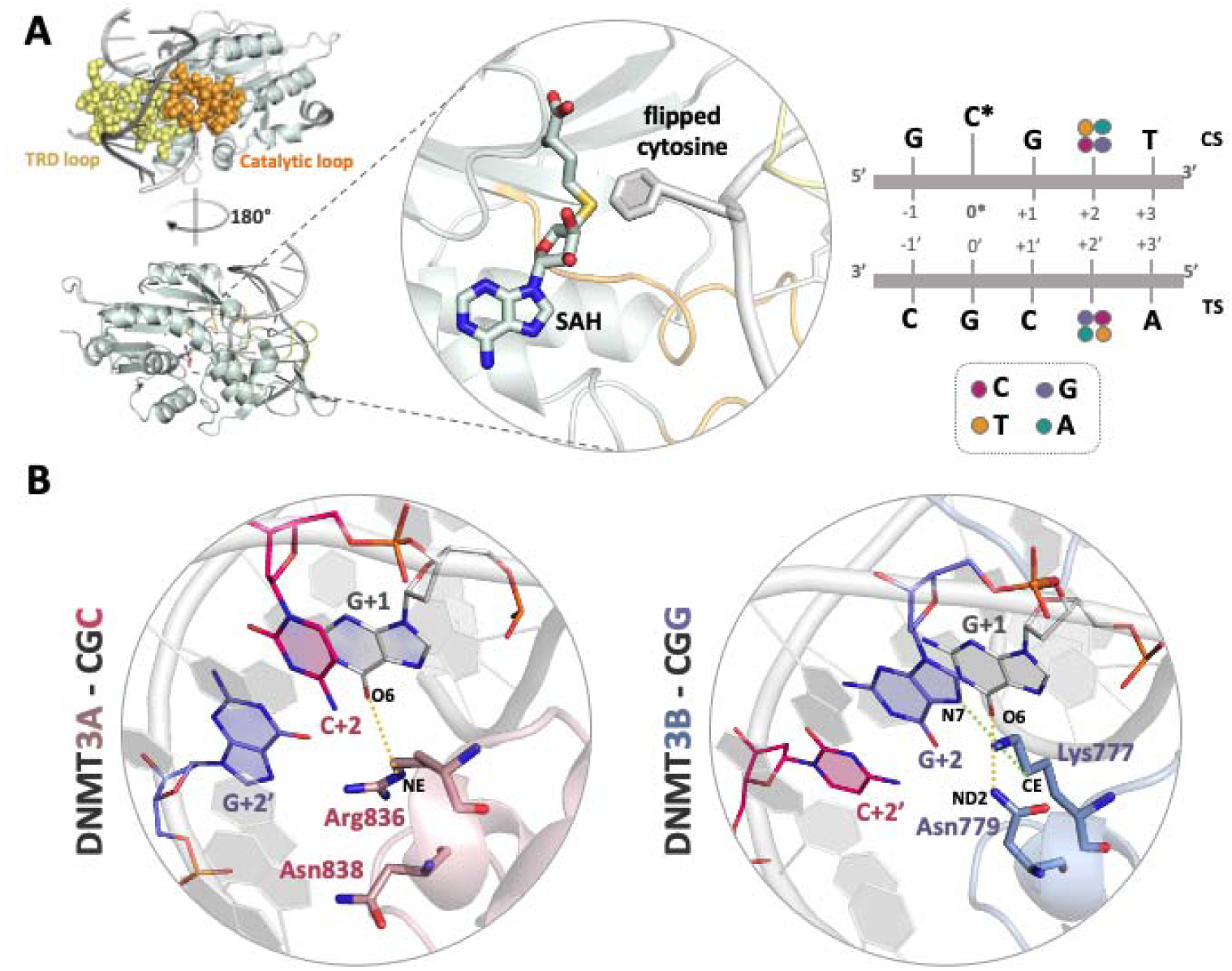
Structural features of DNMT3A and DNMT3B MTase domains. **(A)** Left: 3D structural depiction of DNMT3A/B-MTase domain bound to its target DNA. Orange spheres represent the catalytic loop, and yellow spheres represent the target recognition domain (TRD) loop. The inset focuses on the SAM methyl donor and the flipped target cytosine. Right: Target DNA site. TS and CS stand for target and complementary strands, respectively. 0* denotes the center of the sequence, which is the cytosine to be methylated. Nucleotide numbers with apostrophe (’) state the complementary strand nucleotides. The nucleotide variations explored in this study fall at the +2 and 2’ positions. **(B)** Crystal structures of cognate human DNMT3A and DNMT3B complexes. Right: DNMT3A is bound to CGC (PDB ID: 6F57^13^). In thi crystal structure, Arg836 forms a base specific hydrogen bond with G+1 of CGC (yellow)^13^). Right: DNMT3B is bound to CGG (PDB ID: 6KDA^10^). In this crystal structure, Lys777 forms van der Waals interactions with G+2 (green)^10^) and Asn779 forms a hydrogen bond with G+1 of CGG (yellow)^14^.

**Figure 2.**
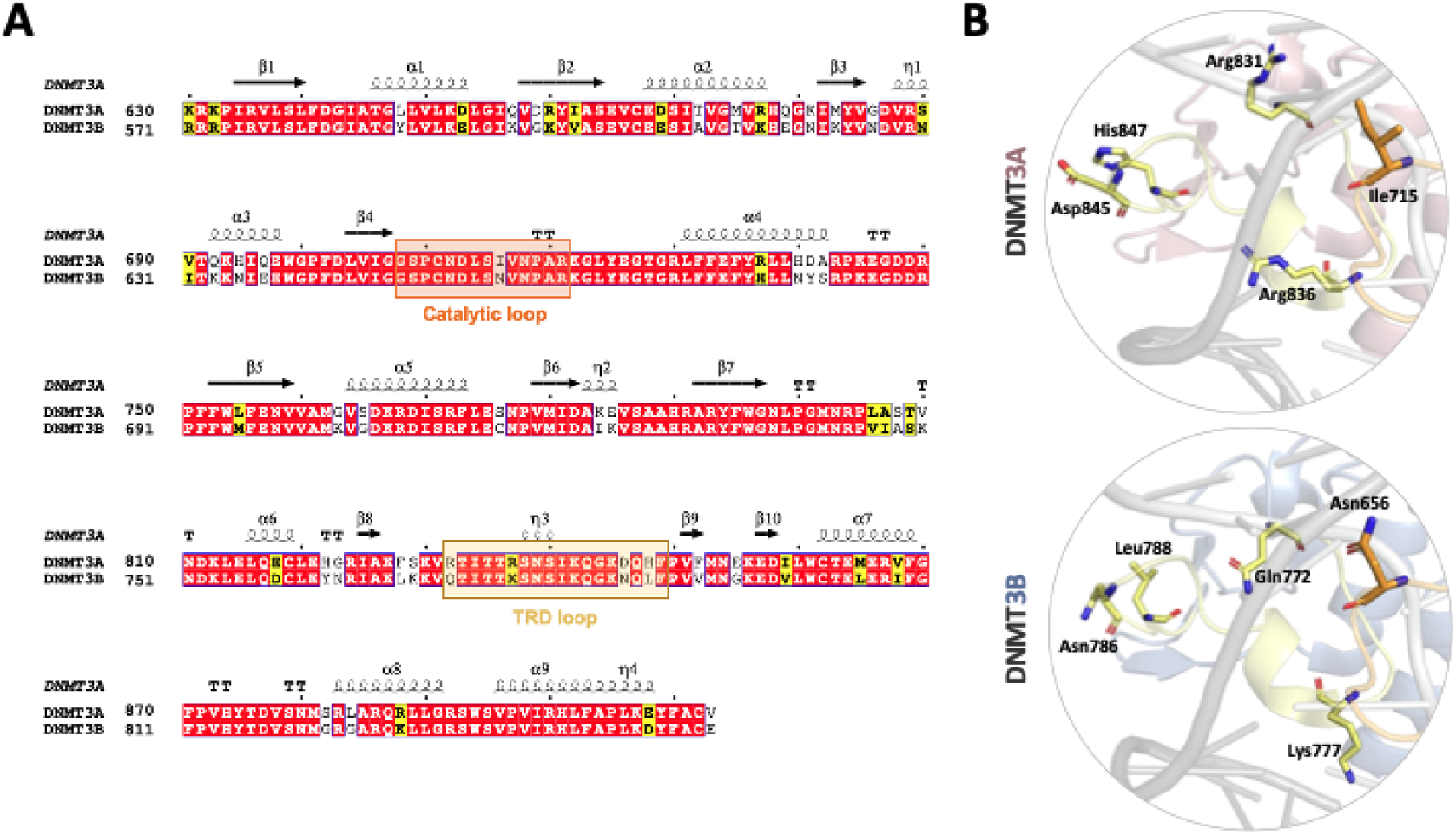
The key differential amino acid positions in DNMT3A and DNMT3B MTase domains. **(A)** A pairwise sequence alignment was performed for the MTase domains of DNMT3A and DNMT3B over their 283 residues. The conserved amino acids are highlighted in red, while the non-conserved ones with similar physicochemical properties are yellow. Black residues with a white background indicate the non-conserved positions. The sequence identity and similarity values are 80.2% and 91.2%, respectively. **(B)** The non-conserved DNMT3A (first row) and DNMT3B (second row) residues located on the major groove contacting TRD-loop (Arg831-to-Gln772, Arg836-to-Lys777, Asp845-to-Asn786, His847-to-Leu788) and on the minor groove contacting catalytic loop (Ile715-to-Asn656) are shown (PDB IDs: 6F57, 6KDA^10,13^). Yellow represents the TRD loop and orange represents the catalytic loop.

Despite their conserved catalytic mechanism, DNMT3A and DNMT3B exhibit distinct DNA sequence preferences. Enzymatic assays and gene knockout studies have shown that DNMT3A preferentially methylates CpG sites followed by a pyrimidine (C or T) at the +2 position (e.g., CGC or CGT), whereas DNMT3B favors purines (G or A) at the same position (e.g., CGG or CGA) ^10,13–16^ (Figure 1A-B). Accordingly, CGC and CGG have been recognized as the primary (i.e., cognate) methylation motifs for DNMT3A and DNMT3B, respectively. The available biochemical experiments have also suggested that DNMT3A and DNMT3B adopt fundamentally distinct methylation strategies: DNMT3A binds DNA in a cooperative and site-specific manner, whereas DNMT3B exhibits processive methylation behavior, scanning along DNA with broader sequence tolerance^17^. In addition to this, deep enzymology studies have demonstrated that the CpG specificity of DNMT3A is higher than DNMT3B within a favored flanking sequence context^14^.

Understanding how these enzymes achieve single-base specificity through distinct methylation mechanisms has remained a major challenge. The specificity has been thought to arise from a small set of residues located within the catalytic loop (residues 707–721 in DNMT3A; 648–662 in DNMT3B) and the target recognition (TRD) loop (residues 831–848 in DNMT3A; 772–789 in DNMT3B)^10,13^ (Figure 2). These loops simultaneously engage the minor groove (catalytic loop) and the major groove (TRD loop) of the DNA, thereby stabilizing the substrate for catalysis ^10,13^ (Figure 1A, 2B). Among all, the structural studies have pinpointed key residues involved in CpG methylation specificity^10,13,14,18^ (Table 1). Among these, Ile715 in DNMT3A and its structurally corresponding residue Asn656 in DNMT3B help shape the catalytic loop conformation through intramolecular interactions, including a hydrogen bond between Asn656 and Arg661 in DNMT3B^14^. Also, the wedge residues Val716 (DNMT3A) and Val657 (DNMT3B) stabilize DNA conformation by occupying the void created during cytosine base flipping, a critical feature for CpG targeting^13,19^. In the TRD loop, the primary guanine readers, Arg836 in DNMT3A and Lys777 in DNMT3B, interact directly with the guanine at the +1 position (G+1). Notably, DNMT3B harbors a secondary guanine reader, Asn779, which further supports sequence selectivity by forming supplementary hydrogen bonds or stabilizing backbone contacts, particularly in contexts where the primary interaction is weakened.

**Table 1.** Functional Roles of Key Residues in DNMT3A and DNMT3B.

| Functional Role | DNMT3A Residue | DNMT3B Residue | Impact on CpG Recognition and Activity |
| --- | --- | --- | --- |
| Wedge Residue | V716 | V657 | Stabilizes DNA conformation by occupying the void created during cytosine base flipping, essential for targeting CpG dinucleotides. Critical for enzyme positioning and engagement with CpG sites.<sup>13,19</sup> |
| Catalytic Loop Stabilizer | I715, R720 | N656, R661 | Stabilizes the catalytic loop via intramolecular interactions (e.g., hydrogen bonding in DNMT3B between N656 and R661).<sup>14</sup> |
| Primary Guanine Reader | R836 | K777 | Directly interacts with the guanine base in CpG dinucleotides, enabling CpG-specific recognition. In DNMT3A, R836 forms key contacts with G+1.<sup>10,13</sup> |
| Secondary Guanine Reader | N838 | N779 | Provides supplementary guanine recognition, stabilizing enzyme-DNA interactions when the primary reader is suboptimal.<sup>10,13</sup> |

While these structural data have provided valuable insights into how DNMT3A and DNMT3B recognize CpG sites through coordinated interactions of catalytic and TRD loops, they do not explain these enzymes’ preferences for the +2 flanking base or their distinct methylation patterns. In 2021, Mallona *et al.* reported that in molecular dynamics (MD) simulations of DNMT3A bound to a CGC sequence, the crystallographically observed Arg836–G+1 interaction was rapidly replaced by an Arg836–G+2′ contact^16^. This rearrangement offers a potential mechanistic explanation for flanking base preferences at the +2 position^16^. This finding highlights the need for further MD simulations to uncover the dynamic determinants of DNMT3 sequence specificity and methylation behavior.

To address this need, we performed a comprehensive MD simulation study of SAM-bound DNMT3A and DNMT3B in complex with four different +2 flanking sequence contexts (CGC, CGT, CGA, and CGG). We also developed a Comparative Dynamics Analysis (CDA) framework to quantify key protein–DNA interaction metrics across a total of 16 µs of all-atom simulations, including hydrogen bonding, electrostatic complementarity, and DNA deformation. We also explicitly tracked the roles of key DNMT3 residues, as outlined in Table 1. Our CDA analyses revealed that DNMT3A achieves high specificity through a rigid, well-aligned interaction network centered on Arg836, favoring compact DNA conformations at CGC motifs. In contrast, DNMT3B discriminates flanking bases through a more distributed and flexible readout involving Lys777 and Asn779, consistent with its processive methylation behavior. Taken together, these findings provide a dynamic explanation for sequence-selective *de novo* methylation and establish a generalizable framework for understanding paralog-specific protein– DNA recognition mechanisms.

## RESULTS AND DISCUSSION

To comprehensively evaluate the sequence selectivity of DNMT3 enzymes, we constructed eight enzyme–DNA complex models based on the available crystal structures: DNMT3A bound to CGC, CGT, CGA, CGG, and DNMT3B bound to CGG, CGT, CGA, CGC. In these models, the DNA sequences differ only at the +2 position, enabling a focused comparison of how protein–DNA interactions are modulated by this single flanking nucleotide (Figure 1A). Each DNMT3 complex was subjected to 2 µs of MD simulations, distributed across four independent 0.5 µs replicates, leading to a total of 16 µs of simulation data. All trajectories were pooled to enhance conformational sampling and improve the statistical robustness of downstream analyses. To isolate sequence-dependent effects and minimize confounding by conserved catalytic features, we excluded interactions involving the SAM cofactor and the flipped cytosine, which are structurally invariant across complexes. Transient contacts observed in fewer than 10% of frames were also filtered out to reduce noise and highlight persistent, functionally relevant interactions. Detailed protocols for model construction and simulation setup are provided in the Methods section.

To systematically quantify protein–DNA interactions across different sequence contexts, we developed a Comparative Dynamics Analysis (CDA) framework that enables parallel evaluation of base and shape readout mechanisms (Figure 3). This framework was specifically designed to capture dynamic and sequence-dependent recognition features that are not accessible from static structures alone. The base readout is defined as the direct recognition of specific nucleobases through hydrogen bonding between polar side chains, such as those of arginine, lysine, and asparagine, and the functional groups exposed in the DNA major groove^20–28^. Therefore, CDA’s base readout module focuses on interactions involving flanking nucleotides at positions +1 to +3 (Figure 3A) and quantifies residue–nucleotide interaction propensities alongside atomistically resolved donor–acceptor geometries across the DNMT3–DNA complexes. The shape readout, in contrast, refers to recognition of DNA geometry and its electrostatic landscape, often mediated by residues such as arginine and lysine that sense changes in the DNA minor groove or phosphate backbone ^22,29–31^. Inspired by previous work demonstrating that features such as minor groove narrowing and DNA backbone deformation can enhance electrostatic complementarity in protein–DNA interactions^24,25,32–36^, we implemented a multi-faceted shape readout module within CDA (Figure 3B). This module first quantifies sequence-dependent deformation using phosphate root-mean-square deviation (P-RMSD) values relative to ideal B-DNA. P-RMSD serves as a dynamic proxy for backbone flexibility. These values are then correlated with local electrostatic interaction energies at the protein–DNA interface to assess how deviations from ideal geometry influence charge complementarity. Additionally, CDA tracks minor groove width as a positional feature and maps residue-level interactions, including hydrogen bonding and hydrophobic contacts, with upstream flanking bases (positions –1, –2, –3), where shape readout is expected to be most influential. To our knowledge, this is the first study to systematically characterize the shape readout in DNMT3 systems.

**Figure 3.**
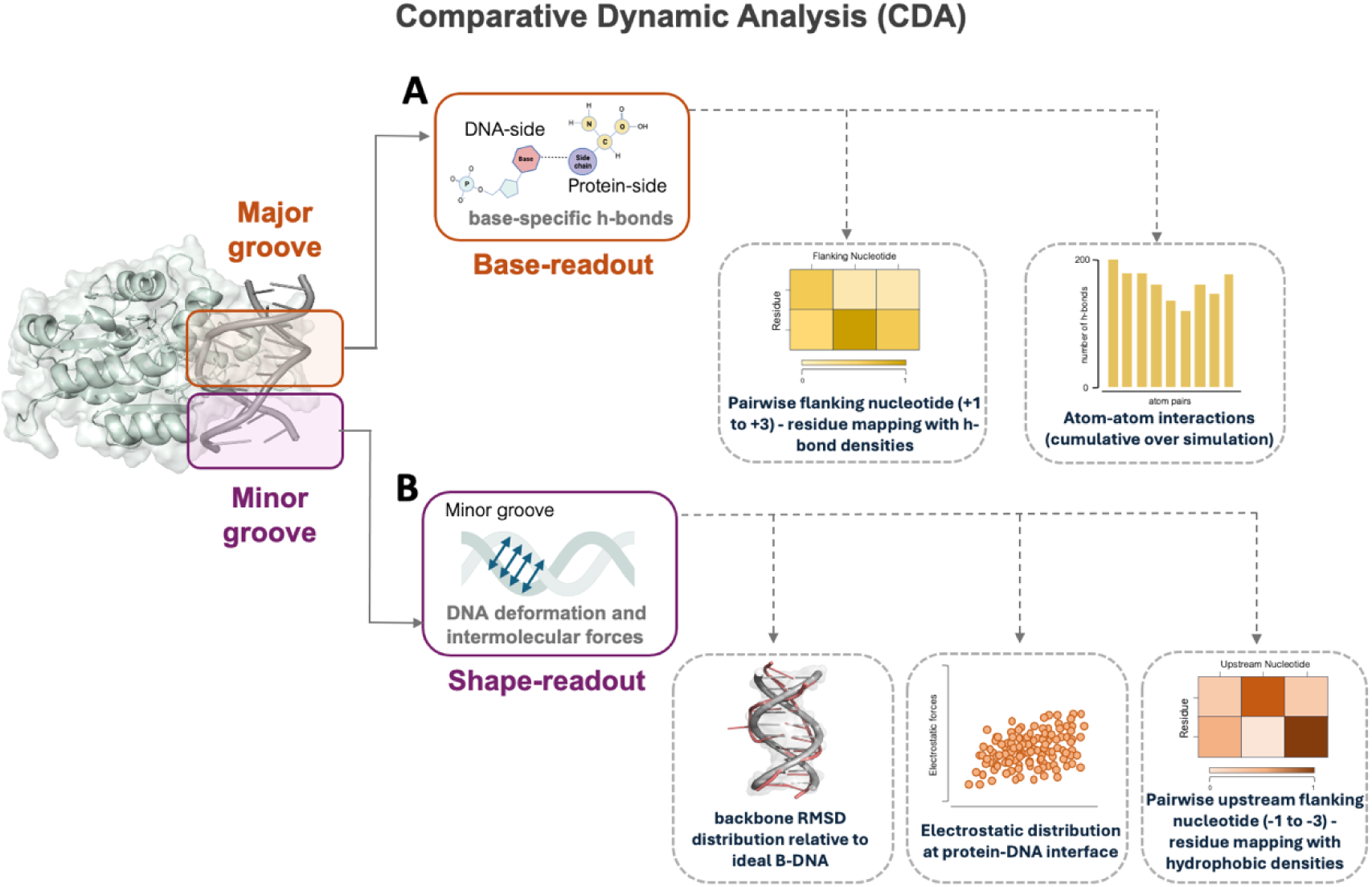
The Comparative Dynamics Analysis (CDA) framework designed to identify DNMT3 base and shape readout rule sets. (A) Base readout. This module quantifies specific hydrogen bond interactions between DNA bases and protein side chains. Step 1: Interaction analysis is performed to detect base-specific hydrogen bonds. The schematic depicts the base-specific hydrogen bonds between DNA base atoms and residue side chain atoms. A heatmap summarizes the average hydrogen bonding intensity between residues and DNA flanking nucleotides at positions +1 to +3. Step 2: The frequency of individual atom-atom hydrogen bonds is visualized through bar plots, representing cumulative interactions over the simulation trajectory. **(B) Shape readout.** This module captures DNA backbone deformation and associated electrostatic forces, especially at the minor groove where shape recognition is most pronounced. Step 1: DNA deformation is quantified using phosphate root-mean-square deviation (P-RMSD) values relative to ideal B-DNA, representing dynamic shape changes. Step 2: These deformations are correlated with electrostatic distributions at the protein–DNA interface, highlighting sequence-dependent complementarity. Step 3: A pairwise interaction matrix maps residue interactions with upstream DNA flanking nucleotides (–1 to –3) based on base-specific hydrogen bonds and hydrophobic contact intensity.

Altogether, our CDA enables an integrated and quantitative analysis of both base-specific recognition and shape-dependent stabilization in a sequence- and residue-resolved manner. Full implementation details are provided in the Methods section. In the following sections, we present key findings from CDA that reveal how DNMT3A and DNMT3B interpret DNA sequence through distinct combinations of base- and shape-readout mechanisms. All CDA scripts and simulation data are publicly available at: https://github.com/CSB-KaracaLab/DNMT3AB_specificity.

### Base readout in DNMT3A is dynamically modulated by the primary guanine rseader Arg836

We began our analysis by applying the base readout module of CDA to DNMT3A–DNA complexes (Figure 3A). This revealed consistent but sequence-dependent patterns of guanine recognition mediated by the primary guanine reader Arg836 (Table 1). In the cognate DNMT3A– CGC complex, Arg836 formed a strong and stable hydrogen bond with the guanine at position +2′ on the complementary strand. In over 70% of simulation frames, this occurred through a bidentate interaction, where two atoms of Arg836’s side chain (NE and NH) simultaneously contacted the N7 and O6 atoms of G+2′ (Figure 4A, Table 2; Supplementary Movie 1). A secondary, less frequent interaction geometry was also observed in ∼20% of frames (Supplementary Figure S1, Table 2).

**Figure 4.**
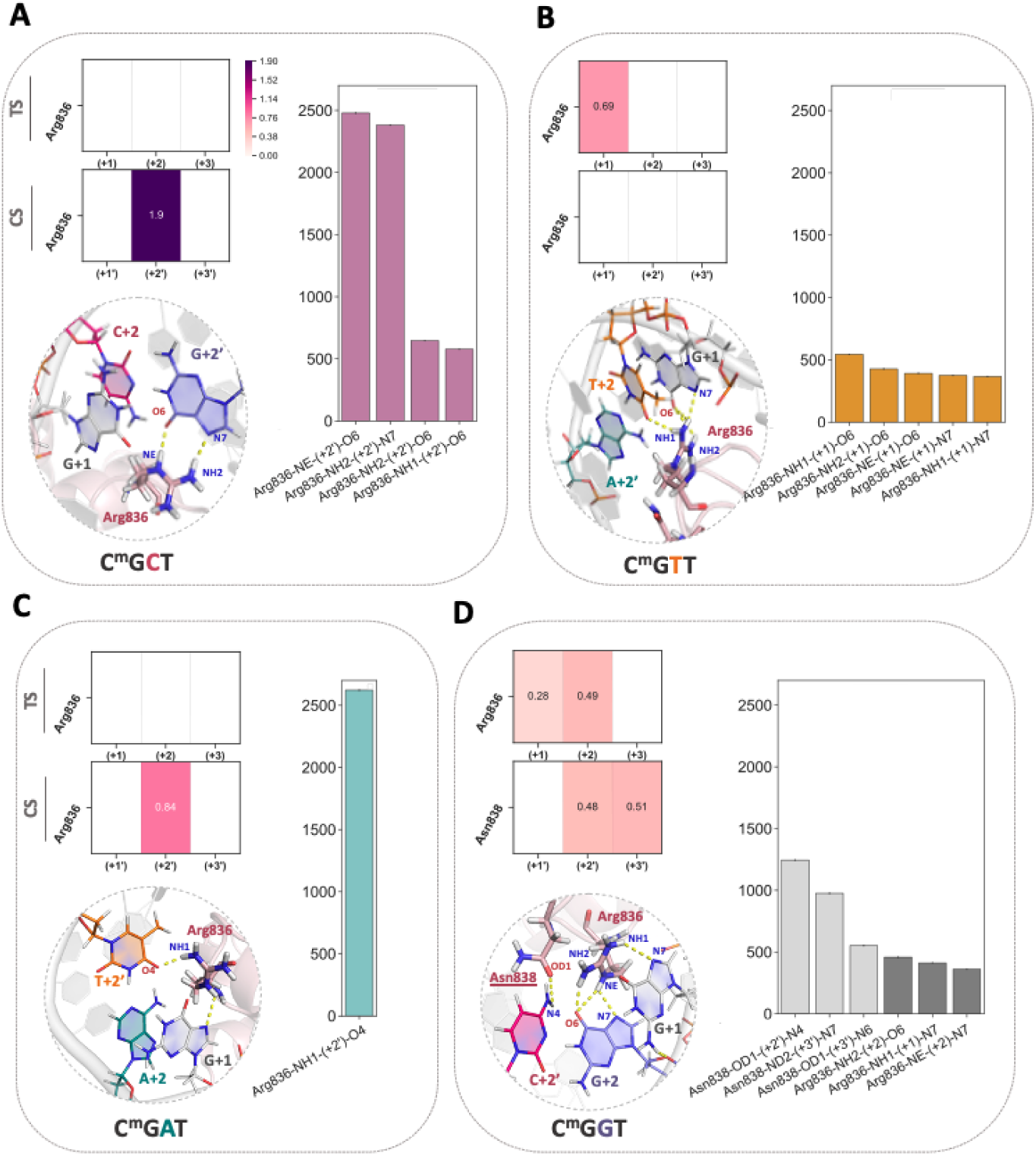
Base readout CDA analysis of DNMT3A-CGX complexes. +1 to +3 flanking nucleotide interactions of DNMT3A with **(A)** CGC, **(B)** CGT, **(C)** CGA, **(D)** CGG sequences. In the heatmaps, the first row represents the target strand (TS), while the second row shows the complementary strand (CS) interactions. The panels contain the distribution of base-specific hydrogen bonds analyzed at the atomistic scale (over 1.6 µs MD trajectory), covering the range of interactions presented in the heatmap. Light colored bars represent the complementary strand interactions, whereas dark colored bars represent target strand interactions. The panels also present the structural snapshots of the important base-specific interactions highlighted in the heatmap and bar plot.

**Table 2.** Summary of base-specific hydrogen bonds formed between DNMT3A with different DNA sequences. The hydrogen bond forming atoms and their observation frequencies are explicitly stated (over the whole 1.6 µs MD trajectory for a given DNMT3A complex).

| DNMT3A | CGC | CGT | CGA | CGG |
| --- | --- | --- | --- | --- |
|  | <u>G+2'</u> | <u>G+1</u> | <u>T+2'</u> | <u>G+2</u> |
| **Arg836** | NE-O6 77.4% | NH1-O6 17% | NH1-O4 81.84% | NH2-O6 14.3% |
|  | NH2-N7 74.3% | NH2-O6 13.3% |  | NE-N7 11.3% |
|  | NH2-O6 20.2% | NE-O6 12.2% |  | <u>G+1</u> |
|  | NH1-O6 18% | NE-N7 11.7% |  | NH1-N7 12.9% |
|  |  | NH1-N7 11.4% |  |  |
| **Asn838** | * | * | * | <u>C+2'</u> |
|  |  |  |  | OD1-N4 38.9% |
|  |  |  |  | <u>A+3'</u> |
|  |  |  |  | ND2-N7 30.5% |
|  |  |  |  | OD1-N6 17.2% |

When the +2 base was changed from C to T in the DNMT3A–CGT complex, this stable Arg836–G+2′ interaction was disrupted. Instead, Arg836 formed weaker and less frequent hydrogen bonds with the guanine at position +1 on the target strand. These interactions involved variable donor atoms and were significantly less stable, resulting in a threefold reduction in interaction intensity compared to the cognate complex (0.69 vs. 1.9; Figure 4B, Table 2). In the DNMT3A–CGA complex, Arg836 partially recovered its binding ability by forming a stable hydrogen bond between its NH1 atom and the O4 atom of T+2′, observed in 82% of conformations (interaction intensity: 0.84; Figure 4C, Table 2). By contrast, in the noncognate DNMT3A–CGG complex, Arg836 was no longer the main contributor to base readout. Instead, the structurally correspondent of DNMT3B’s secondary guanine reader, Asn838, took a more prominent role. In the DNMT3A–CGG complex, Asn838 formed a hydrogen bond with the N4 atom of C+2′ in 39% of frames (Figure 4D, Table 2). Meanwhile, Arg836 exhibited only weak and infrequent interactions with G+1 and G+2 (interaction intensities: 0.28 and 0.49), and bidentate geometries were rare.

These findings extend previous structural observations. In the CGC context, our simulations identified a stable Arg836–G+2′ bidentate interaction that was not reported in the work of Mallona *et al.*. This mode of stable guanine recognition aligns with the prior work underscoring that arginine commonly forms multi-point hydrogen bonds with guanine bases in the major groove, a well-established strategy among DNA-binding proteins^22,28^. In the context of other sequences, the available structures show that Arg836 contacts G+1 in CGT and A+2 in CGA (Figure 1B) (PDB IDs: 6F57^13^, 5YX2^13^, 6W8B^19^). In consistent with these structural data, we observed stable Arg836–G+1 interactions in both CGT and CGA, but also identified additional geometries and donor–acceptor pairings that emerged only in dynamics. Finally, our results also suggest a backup reader role for Asn838 in the context of CGG.

### DNMT3B’s base readout is mediated by the cooperative action of its primary Lys777 and secondary Asn779 guanine readers

We applied the base readout module of CDA to DNMT3B–DNA complexes too. As an outcome, we observed that in the cognate DNMT3B–CGG complex, the primary guanine reader Lys777 formed moderate- to low-intensity hydrogen bonds with bases on the target strand, namely G+1, G+2, and T+3, with interaction intensities ranging from 0.43 to 0.17. Here, Lys777 did not contact the bases on the complementary strand (Figure 5A). Instead, the secondary guanine reader Asn779 played a key role, forming hydrogen bonds with both G+1 and C+2′ with an average interaction intensity of ∼0.38. The most frequent interactions observed were between Lys777 NZ and G+1 N7 (37.5% of conformations), and between Asn779 OD1 and C+2′ N4 (30.1%) (Table 3). Notably, this distributed recognition pattern mirrors the cooperative readout mode seen in the DNMT3A–CGG complex, suggesting that certain DNA sequences may impose preferred modes of protein engagement.

**Figure 5.**
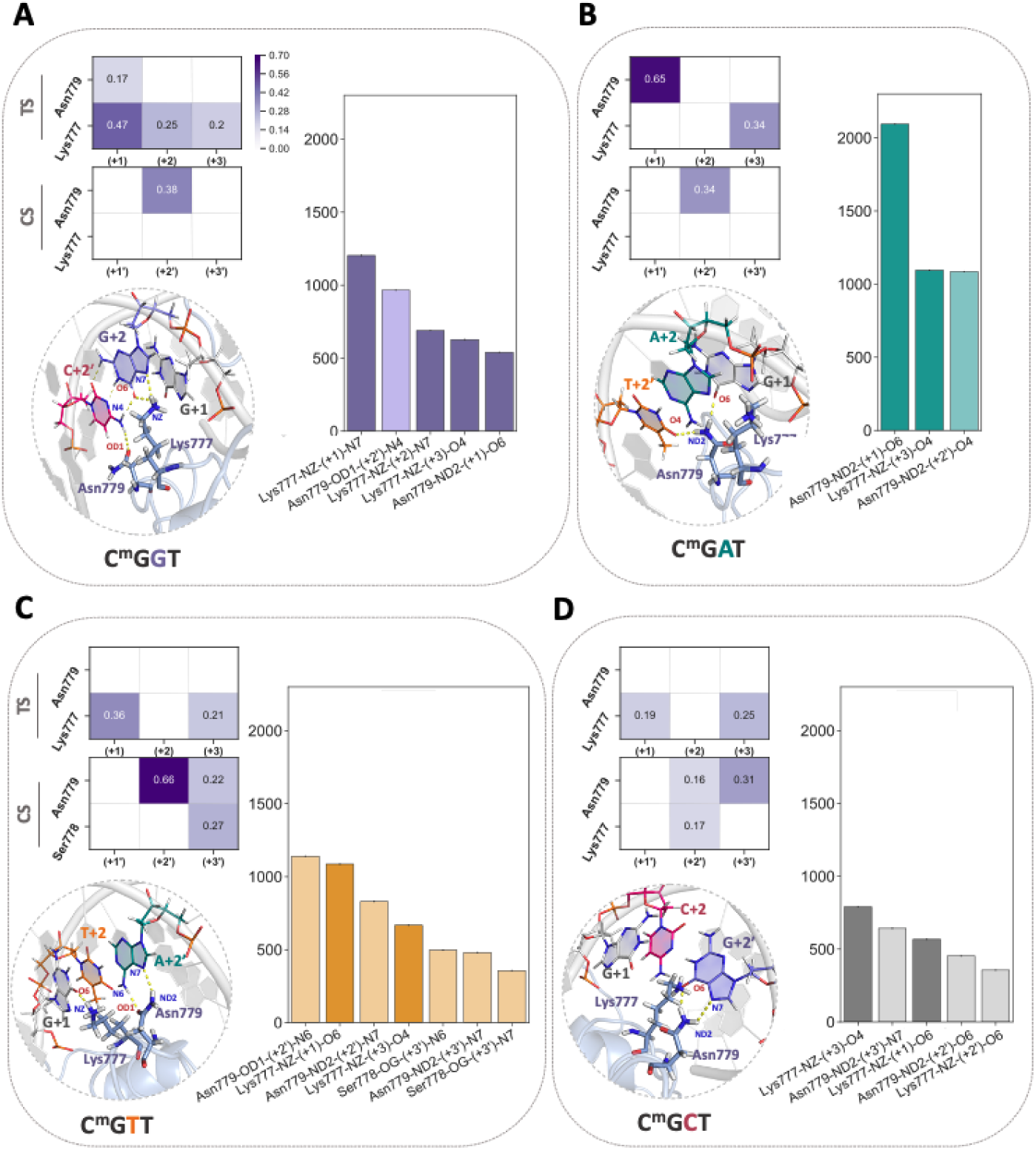
Base readout CDA analysis of DNMT3B-CGX complexes. +1 to +3 flanking nucleotide interactions of DNMT3A with **(A)** CGC, **(B)** CGT, **(C)** CGA, **(D)** CGG sequences. In the heatmaps, the first row represents the target strand (TS) interactions, while the second row shows the complementary strand (CS) ones. The panels contain the distribution of base-specific hydrogen bonds analyzed at the atomisti scale (over 1.6 µs MD trajectory), covering the range of interactions depicted by the heatmap. Light colored bars represent the complementary strand interactions, whereas dark colored bars represent target strand interactions. The panels also present the structural snapshots of the important base-specifi interactions highlighted in the heatmap and barplot.

**Table 3.**
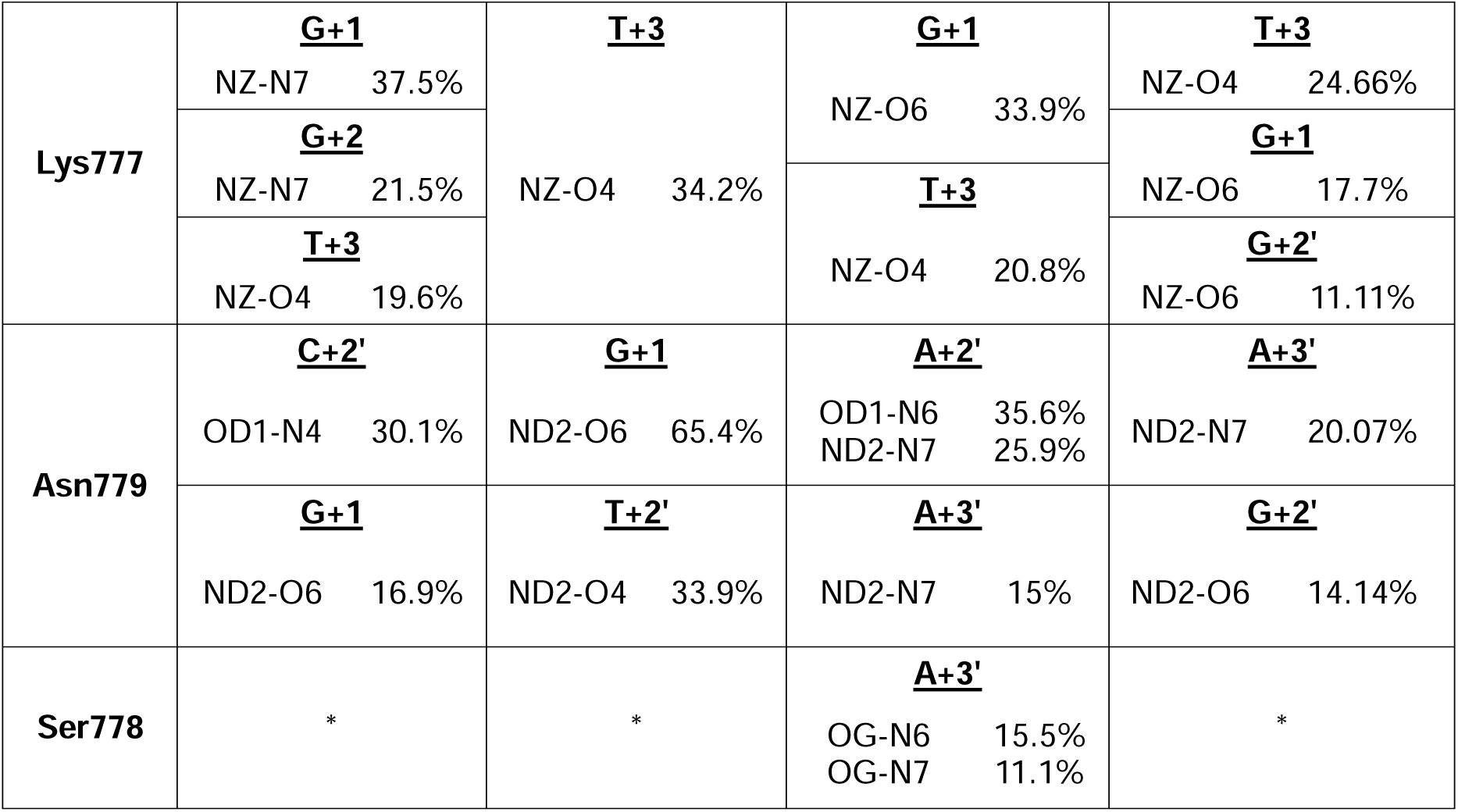
Summary of base-specific hydrogen bonds formed between DNMT3B and different DNA sequences. The hydrogen bond forming atoms and their observation frequencies are explicitly stated (over the whole 1.6 µs MD trajectory for a given DNMT3A complex).

In the DNMT3B–CGA complex, the recognition landscape shifted. Lys777 redirected its interactions toward T+3 (intensity: 0.34), while Asn779 retained strong engagement with G+1 (intensity: 0.65). It also contacted T+2′ with a similar frequency to that one observed in CGG (∼34%) (Figure 5B, Table 3). In the CGT complex, Asn779 exhibited its strongest base-specific engagement, forming a bidentate hydrogen bond with A+2′ through its ND2 and backbone atoms (intensity: 0.66), alongside secondary contacts with A+3′ (intensity: 0.22). These interactions leveraged adenine’s dual donor–acceptor capacity (Supplementary Figure S3, Figure 5C). The adjacent Ser778 contributed to recognition by forming a hydrogen bond with A+3′ via its OG atom. In this sequence, Lys777 remained in contact with G+1, which dynamically alternated between G+1, T+3, and G+4′, displaying a dual binding mode (Supplementary Figure S2). By contrast, the DNMT3B–CGC complex exhibited a weaker and more dispersed interaction pattern. In this case, Lys777 formed only low-intensity hydrogen bonds with G+1 and T+3 (intensities: 0.19–0.25), and occasional contacts with G+2′ (0.17). In DNMT3B–CGC complex the interaction between Asn779 and +2′ base dropped more than fourfold compared to CGT, shifting toward transient engagements with G+2′ and A+3′ (Figure 5D). Interestingly, Lys777 formed a stable hydrogen bond with G+4′ O6 in 75% of conformations. In the remaining 25%, it changed between G+1 and G+2′, further underscoring DNMT3B’s flexible and lower-specificity recognition mode in the absence of a favorable +2 base (Supplementary Figure S2).

Comparison of these results with available crystal structures revealed both agreements and new insights. The available structural data suggest that Lys777 forms van der Waals contacts with G+2 in CGG and with T+2 or A+2 in CGT and CGA, respectively (PDB IDs: 6KDA^10^, 6KDB^10^, 6U8P^14^). Across all complexes, Asn779 was shown to interact with G+1. Our simulations corroborate these findings and further demonstrate that the Lys777–Asn779 network adapts to local sequence, forming distinct hydrogen bonding geometries depending on the +2 base. In particular, the high-occupancy bidentate interaction involving Asn779 in DNMT3B–CGT, which is absent in crystal structures, reveals a dynamic mechanism of base-specific recognition not previously observed. Also, our simulations recapitulate the Asn779–T+2′ contact observed in DNMT3B–CGA^14^ and identify a new interaction in which Lys777 forms a hydrogen bond with the O4 atom of T+3 in 34.2% of conformations (Figure 5B).

Also, within the perspective of base readout, considering with the denser and more stable base-specific hydrogen bonding profiles observed for DNMT3A, our findings strikingly help to explain Gao *et al.*’s following observation^14^: DNMT3A exhibits higher CpG specificity than DNMT3B within its favored sequence contexts. Simply put, while DNMT3B employs a flexible and cooperative recognition network that accommodates a broader range of sequences, DNMT3A relies on more rigid and focused base-specific contacts, consistent with its sharper sequence preference profile.

### DNMT3A shape readout is highly sensitive to +2 flanking nucleotide identity

To assess how DNA shape contributes to sequence specificity in DNMT3A, we applied the shape readout module of our CDA framework. Using P-RMSD as a proxy for sequence-dependent backbone flexibility, we found that the DNMT3A–CGC complex exhibited the broadest and most deformable backbone profile, with P-RMSD values ranging from 1.5 Å to 4.5 Å (Supplementary Figure S4A). This wide distribution reflects both compact base-pairing and local flexibility, consistent with CGC’s optimal fit within the DNMT3A binding pocket. In contrast, CGA displayed the narrowest deformation profile. These differences in DNA deformability were closely tied to electrostatic complementarity. Among all sequence contexts, CGC exhibited the strongest positive correlation between P-RMSD and electrostatic energy (r = 0.57), indicating that as the DNA backbone deviates from ideal B-DNA, electrostatic matching at the protein– DNA interface is gradually lost. CGC also showed the lowest total electrostatic interaction energies, reinforcing its role as the preferred flanking sequence. In contrast, CGT and CGG displayed weaker correlations (r ∼ 0.20), and CGA showed no significant relationship (Figure 6A-B). Minor groove width analysis supported these findings: all sequences showed more compact grooves relative to the reference crystal structure, but CGC consistently exhibited the narrowest groove at higher nucleotide indices. Notably, CGT closely resembled CGC in this regard (Supplementary Figure S4B).

**Figure 6.**
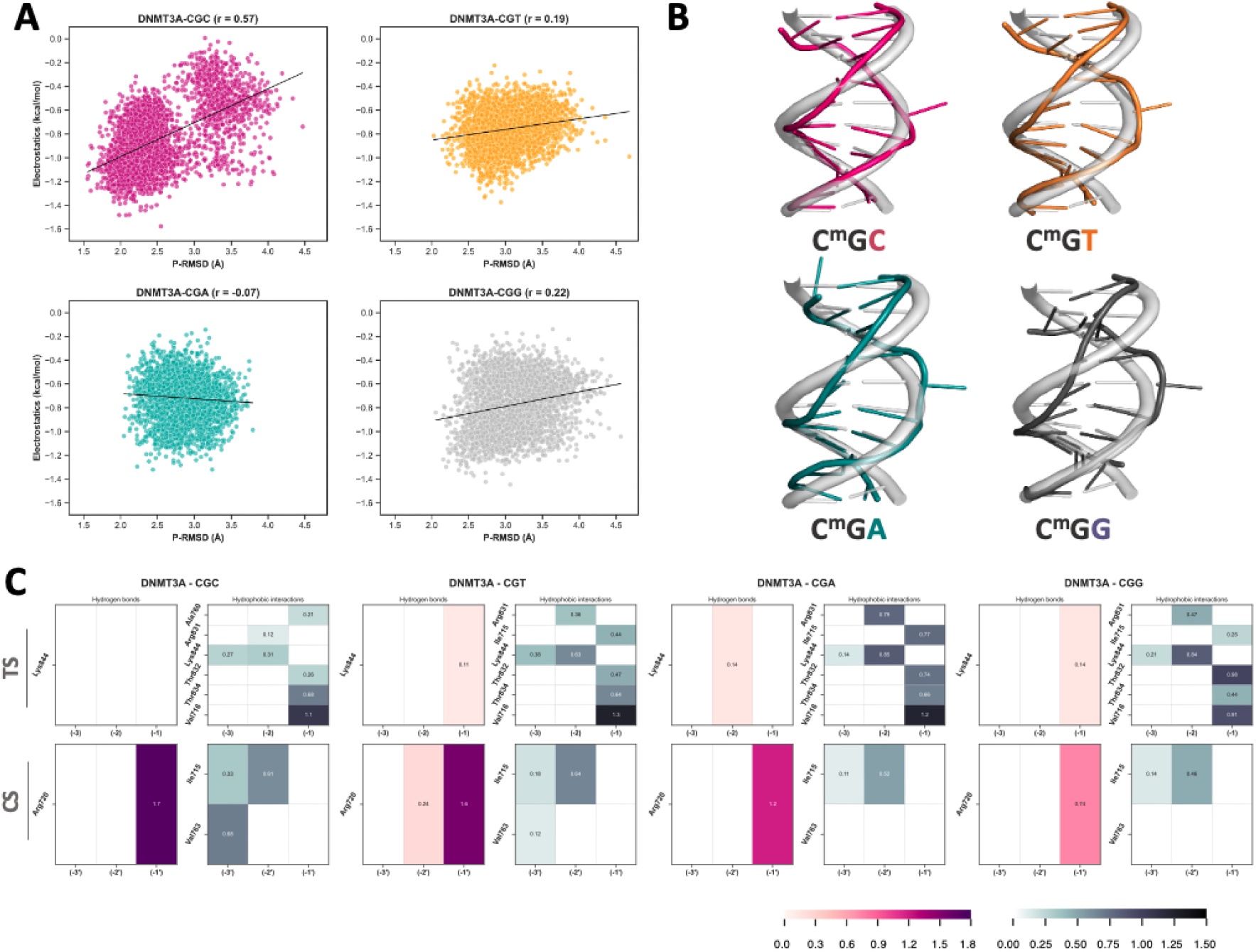
Shape readout CDA analysis of DNMT3A–CGX complexes. Sequence-dependent DNA deformation and electrostatic interactions for DNMT3A bound to CGC, CGT, CGA, and CGG sequences. **(A)** Scatter plots show correlation between phosphate backbone RMSD (P-RMSD, Å) relative to ideal B-DNA and electrostatic interaction energies (kcal/mol) at the DNA–protein interface, calculated across the 2 µs MD simulations. Pearson correlation coefficients (r) indicate the strength of coupling between structural deformation and electrostatic complementarity. **(B)** Representative structural snapshots of DNA backbone conformations illustrate sequence-specific minor groove deformation. DNA strands are colored according to the sequence context, emphasizing deformation around the methylated CpG site. **(C)** Interaction heatmaps quantify residue-level shape recognition features, including hydrogen bonds and hydrophobic contacts at positions –1 to –3. Top rows depict target strand (TS) interactions, and bottom rows show complementary strand (CS) interactions.

To uncover the molecular basis for these shape-dependent effects, we analyzed residue-level hydrogen bonding and hydrophobic interactions at upstream flanking positions (–1 to –3), where shape readout is expected to be most influential (Figure 6C). The catalytic loop stabilizer Arg720 formed base-specific hydrogen bonds with G(–1) in all complexes, with interaction frequencies strikingly reflecting DNMT3A’s methylation preference (CGC > CGT > CGA > CGG). In the CGC complex, the wedge residue Val716 formed a strong hydrogen bond with G(–1) (intensity: 1.1), while the catalytic loop stabilizer Ile715 contributed significant hydrophobic contacts with the –2′ and –3′ bases (intensities: 0.33 and 0.61), helping anchor the complementary strand in the minor groove. Additional stabilizing interactions involved the TRD Thr834, Lys844, and Val763 at –3′. In the CGT complex, Arg720 maintained high-frequency hydrogen bonding at –1′ and –2′, while Val716 and Ile715 retained moderate interaction levels. CGA showed increased hydrophobic engagement on the target strand, but reduced participation from Ile715 and Arg720. The CGG complex exhibited the weakest interaction network overall, with reduced Arg720 hydrogen bonding, minimal Ile715-mediated hydrophobic stabilization, and loss of Val763 engagement. In contrast, TRD residues Thr832 and Lys844 slightly increased their contacts in CGG, potentially compensating for weakened catalytic loop interactions (Figure 6C).

Together, these results illustrate how DNMT3A integrates base and shape readout mechanisms to discriminate flanking sequences. In the CGC complex, both mechanisms are maximally engaged: Arg836 forms a stable bidentate bond with G+2′, while shape readout residues, particularly Arg720, Val716, and Ile715, stabilize a flexible and electrostatically optimized DNA conformation. In CGT, base readout is weaker, but the shape remains favorable, allowing intermediate recognition. In CGA, base readout is strong, but the DNA is rigid and poorly matched to the protein surface, limiting shape readout and lowering overall preference. Finally, CGG lacks both strong base contacts and shape compatibility, resulting in the weakest recognition. The central role of Arg720 in anchoring deformable upstream bases across all favored sequences further highlights how dynamic shape sensing complements base-specific interactions to define DNMT3A sequence selectivity.

### Ile-to-Asn catalytic loop substitution dictates DNMT3B shape readout

To extend our shape readout analysis to DNMT3B, we assessed the DNA deformation across all DNMT3B–DNA complexes by calculating P-RMSD values relative to ideal B-DNA. Unlike DNMT3A, where deformation varies significantly between motifs, nearly all DNMT3B complexes achieved low P-RMSD values (∼1.5 Å), suggesting generally higher tolerance for structural flexibility (Supplementary Figure S4C). Among these, the cognate CGG motif exhibited the widest P-RMSD range (1.5–4.5 Å), while the non-cognate CGC complex showed the narrowest distribution (up to 3.5 Å). We next examined how these structural variations affect protein–DNA energetics. In DNMT3B–CGG, P-RMSD values are positively correlated with electrostatic energies (r = 0.46), as in the cognate DNMT3A complex (Figure 7A). CGA showed a weaker positive trend (r = 0.17), while CGT exhibited a mild negative correlation (r = –0.23). CGC displayed negligible correlation, suggesting limited sensitivity to shape variation. Minor groove analysis showed that, as in DNMT3A, all DNMT3B complexes exhibited narrower grooves than their crystal structure counterparts. However, unlike DNMT3A, the profiles were nearly identical across all sequences (Supplementary Figure S4D, Figure 7B).

**Figure 7.**
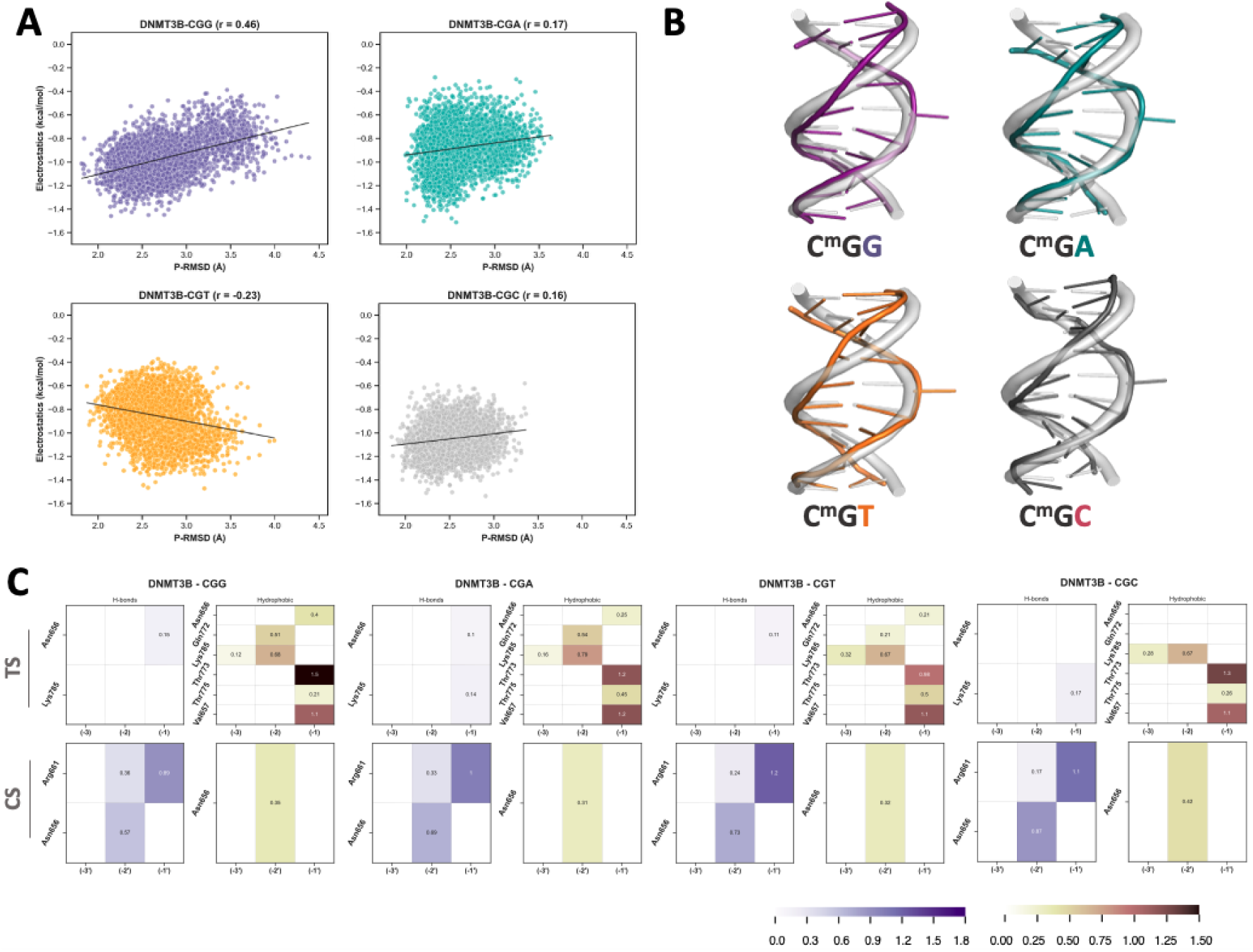
Shape readout CDA analysis of DNMT3B–CGX complexes. Sequence-dependent DNA deformation and electrostatic interactions for DNMT3B bound to CGG, CGA, CGT, and CGC sequences. **(A)** Scatter plots show correlation between phosphate backbone RMSD (P-RMSD, Å) relative to ideal B-DNA and electrostatic interaction energies (kcal/mol) at the DNA–protein interface, calculated across the 2 µs MD simulations. Pearson correlation coefficients (r) indicate the strength of coupling between structural deformation and electrostatic complementarity. **(B)** Representative structural snapshots of DNA backbone conformations illustrate sequence-specific minor groove deformation. DNA strands are colored according to the sequence context, emphasizing deformation around the methylated CpG site. **(C)** Interaction heatmaps quantify residue-level shape recognition features, including hydrogen bonds and hydrophobic contacts at positions –1 to –3. Top rows depict target strand (TS) interactions, and bottom rows show complementary strand (CS) interactions.

To identify the molecular basis for these observations, we analyzed residue–nucleotide interactions at upstream positions (–1, –2, –3). The catalytic loop stabilizer Arg661, the DNMT3B counterpart of DNMT3A’s Arg720, consistently formed hydrogen bonds with bases at –1′ and –2′ across all motifs, with highest occupancy in CGT and CGC (Figure 7C). The non-conserved substitution Asn656, replacing DNMT3A’s Ile715, played a dual role by forming both hydrogen bonds and hydrophobic contacts, particularly with the –2′ base. The wedge residue Val657 and Thr773 consistently formed hydrophobic contacts with G(–1), while Lys785 and Thr775 contributed moderately at upstream sites, especially in CGA and CGT. In CGG, Asn656 formed multiple hydrogen bonds at –1′ and –2′, while Thr773 and Val657 stably engaged G(–1). In CGA and CGT, Arg661 and Asn656 maintained moderate interactions, with Thr773 and Lys785 contributing at –1 and –2. In CGC, Asn656 retained contacts at –2′, but target strand interactions and overall hydrophobic contributions were reduced. Nonetheless, Thr773 and Val657 continued to interact with G(–1), maintaining minimal shape anchoring (Figure 7C).

These results help explain DNMT3B’s flanking sequence preference order (CGG > CGA > CGT > CGC) as a product of complementary base and shape readout. In CGG, the primary and secondary guanine readers (Lys777 and Asn779) are most effectively engaged, while upstream residues, particularly Asn656, Arg661, Thr773, and Val657, collectively support shape stabilization. In CGA, base readout is weaker, but shape anchoring remains strong, sustaining moderate recognition. In CGT, base interactions decline further, though some shape-readout contributions persist. Finally, in CGC, both base and shape readout are diminished: target strand interactions are minimal, and upstream stabilizers are less engaged. These observations highlight DNMT3B’s reliance on a flexible and distributed readout network, capable of accommodating multiple sequence contexts but favoring those where moderate base recognition aligns with consistent, if shallow, shape engagement.

### Distinct Methylation Strategies Are Grounded in Electrostatics and Flexibility

As outlined by our CDA analysis, compared to DNMT3B, DNMT3A forms denser base-specific interactions and exerts tighter control over DNA shape, while DNMT3B exhibits weaker sequence-specific recognition and limited shape sensitivity. To further explore the distinct methylation strategies proposed by Norvil et al.^17^, cooperative and site-specific for DNMT3A, processive and sequence-tolerant for DNMT3B, we extended our analysis to include ionic interactions and loop flexibility analysis.

Our ionic interaction analysis demonstrated that DNMT3A–CGC forms strong electrostatic anchoring contacts through TRD loop residues such as Arg831 and Lys844, reinforcing a highly stable and specific protein–DNA interface. Unlike DNMT3A, DNMT3B lacks strong ionic engagement via TRD loop residues due to the Arg831-to-Gln772 substitution, resulting in reduced electrostatic stabilization. In DNMT3B–CGG, ionic interactions were more distributed and primarily involved non-TRD residues such as Lys782 and Arg823, supporting a looser and more flexible engagement mode (Supplementary Figure S5). RMSF analysis further revealed that DNMT3A’s catalytic loop is inherently more rigid in the unbound state and becomes further stabilized upon DNA binding, consistent with a pre-organized, site-specific recognition mechanism. In contrast, DNMT3B’s catalytic loop remains comparatively dynamic even in the bound state, supporting a scanning, processive engagement strategy (Supplementary Figure S6-7). Notably, the TRD loop is more flexible in DNMT3A than in DNMT3B, indicating that while DNMT3A depends on rigid catalytic loop anchoring, it may use TRD flexibility to fine-tune sequence discrimination (Supplementary Figure S8).

Together, these results show that DNMT3A couples strong base and shape readout with tight electrostatic anchoring, whereas DNMT3B relies on flexible loops and weaker, distributed contacts to accommodate diverse flanking sequences. This mechanistic distinction provides a structural explanation for the divergent methylation behaviors originally proposed by Norvil et al.^17^

## CONCLUSION AND OUTLOOK

DNMT3A and DNMT3B share highly similar catalytic domains with 91% sequence similarity. Yet they exhibit distinct preferences for CpG flanking sequences and adopt divergent methylation strategies. Using our CDA approach, we found that DNMT3A achieves high specificity through a rigid, sequence-specific hydrogen bonding network and strong TRD-mediated electrostatic anchoring. In contrast, DNMT3B utilizes a more flexible and distributed recognition strategy, enabling broader substrate tolerance consistent with its processive methylation behavior.

To achieve this, we quantified, for the first time, the relative contributions of base and shape readout mechanisms across different CGX sequence contexts. Although we did not compute absolute binding free energies, our CDA framework allowed us to extract residue-level, sequence-specific interaction patterns that reveal how recognition is encoded at the atomistic level. Also, for focusing on how flanking sequences influence the stabilization of the protein– DNA interface, we modeled SAM-bound, pre-reactive complexes, isolating recognition mechanisms from catalytic events. A complete mechanistic understanding of catalysis in different sequence contexts would require hybrid quantum mechanics/molecular mechanics simulations, which fall outside the scope of this study but represent an important avenue for future work.

To ensure robustness, we performed all simulations in quadruplicate. Most key base-specific interactions were reproducible across replicates, while variability in lower-frequency contacts reflected inherent conformational flexibility. Pooling data across runs allowed us to emphasize consistent, functionally relevant features while minimizing noise from rare or transient events. Furthermore, we acknowledge the stabilizing role of non-sequence-specific interactions, such as backbone contacts, which are less sensitive to nucleotide identity. These were deliberately excluded from base readout analyses to maintain a clear focus on sequence-discriminative features.

Upon integrating our simulation outcomes with prior biochemical studies^14–16,27,38^, we bridged experimentally observed patterns of flanking sequence preferences and methylation behavior with the underlying atomistic-level interactions. We achieved this by focusing on core CpG contexts. Our reductionist modeling strategy was a intentional design choice, in line with Occam’s razor, relying on favoring the simplest system capable of explaining the observed specificity. However, we are also aware that the emerging evidence suggests that specificity extends to broader flanking regions (e.g., –3 to +3), non-CpG substrates, and potential allosteric modulation by cofactors such as DNMT3L^14,38–40^. Future work should expand CDA to explore these dimensions.

Finally, this study builds on our previous efforts^37,41,42^ to map dynamic recognition at biomolecular interfaces. It underscores the multifactorial nature of sequence specificity, where recognition arises from the interplay of base contacts, DNA shape, and interfacial electrostatics, rather than from any single dominant interaction. Looking ahead, as future work, we envision to combine the atomistic resolution of MD with the generalization capacity of machine learning, using CDA-derived features to predict recognition rules and specificity-determining positions across protein families.

All in all, our study provides a dynamic, generalizable framework for understanding and engineering protein–DNA recognition at base-pair resolution, laying the groundwork for future advances in targeted DNA methylation and the rational reprogramming of paralogous systems.

## METHODS

### Pairwise sequence alignment

Pairwise sequence alignment of MTase domains of DNMT3A and DNMT3B was performed on the EMBOSS Needle web-server^43^. Default settings and the EBLOSUM62 matrix were used for alignment. Alignment results were visualized using Espript 3.0 web-server^44^.

### Molecular dynamics simulations

To assess the sequence selectivity of DNMT3A and DNMT3B enzymes, all-atom molecular dynamics simulations in explicit solvent were performed to capture atomistic interactions between protein and DNA. Atomic coordinates of DNMT3A-DNMT3L complex with single CpG-containing DNA (PDB ID: 6F57^13^) and the human DNMT3B-DNMT3L complex with DNA containing a CpGpG site (PDB ID: 6KDA^10^) were used as starting structures for MD simulations. We started residue numbering from 1 to allow for direct sequence-based comparison between DNMT3A and DNMT3B constructs. 10 bp long CGC-containing DNA (5’-CATG*CG**C**TCT-3’) of DNMT3A-CGC complex were used as template DNA. The cytosine at +2 in DNMT3A-CGC modified to the other three nucleotide possibilities, i.e., thymine, adenine, and guanine using 3DNA web server^45^. The modeled DNA structures were further isolated to generate the corresponding DNMT3B complexes. To create the pre-reaction forms of these complexes, the SAH molecule was replaced by the S-Adenosylmethionine (SAM) molecule in the crystal structure of mouse DNA methyltransferase 1 with AdoMet (PDB ID: 3AV6^46^). Additionally, zebularine in the crystal structure of DNA was converted to cytosine. All created DNMT3A-CGC/T/A/G and DNMT3B-CGC/T/A/G complexes were refined with the advanced refinement protocol using the HADDOCK2.2 web-server Guru Interface, and 200 structures were generated for each system^47^. The structures with the best scores were selected for molecular dynamics simulations. For DNA-unbound simulations, DNA molecules were removed from the refined structures.

MD simulations were performed using GROMACS 2020.4^48^ under the effect of AMBER14SB-parmbsc1^49^ force field. Parameters of SAM cofactor were generated by the Acpype web-server^50^. In the initial structures, the 5′ and 3′ DNA ends were capped to prevent nonnatural electrostatic effects. The simulation box was set up as a rhombic dodecahedron with a minimum distance between the solute and the edge of the box of 1.4 nm. Before running the MD simulations, the force constant of the positional restraints were increased from 1000 to 2500 kJ mol^-1^ nm^-2^ and protein-DNA complexes were minimized by the steepest descent algorithm in the vacuum. Then, they were solvated with TIP3P water^51^ and 0.15 M KCl and the system was relaxed with energy minimization again.

The systems were heated to 310 K using the velocity-rescaling thermostat^52^. NVT equilibration was completed in 6 steps, each lasting 20 ps, with each step after the first step force constants of the positional restraints were decreased from 2500 to 500, 500 to 400, 400 to 300, 300 to 200, and 200 to 100 kJ mol^-1^ nm^-2^. Then sequential 20 ps MD simulations with gradual release of position restraints on heavy atoms were performed under constant pressure at 1 bar using Berendsen barostat^53^. Before starting the MD run, the positional restraints were completely removed. For each DNA-bound system, 500 ns length 4 independent simulations were performed with different initial velocities (total of 2 µs for each system). DNA-unbound systems were simulated for 500 ns. The PME approach^54^ was employed to address long-range electrostatic interactions, utilizing a nonbonded cutoff of 1.2 nm. The LINCS method^55^ was used to restrict hydrogen bond lengths. After accounting for periodic boundary conditions, an ensemble was created by taking 1 frame every 0.5 ns for each simulation. Before the analysis part, equilibration time was set as 100 ns and discarded from the simulations. As quality control measures, we calculated RMSD and RMSF values of each complex with GROMACS 2020.4 software^48^ (Supplementary Figures S6-9). For clarity and consistency with external references, in the analysis part we adjusted the numbering to match UniProt annotations by adding 629 for DNMT3A and 570 for DNMT3B, and we use these UniProt-based residue numbers throughout the manuscript.

### Calculation and analysis of the non-covalent interactions

Intra- and intermolecular interactions were calculated using the Interfacea Python package^56^, enabling quantification of non-covalent contacts including hydrogen bonds, salt bridges, and hydrophobic interactions. Significant interactions at residue-level and atom-level were defined as those present in at least 10% of the simulation frames. Electrostatic interaction energies between protein and DNA were computed with the FoldX suite^57^ using the AnalyseComplex function and the complexWithDNA flag. Outputs from Interfacea and FoldX for each simulation frame were processed using Bash scripting and the Pandas library^58^ in Python. Statistical visualizations of interaction distributions were generated using the Seaborn library^59^.

### DNA geometry analysis

To derive DNA geometrical features, we calculated the minor groove width and phosphate backbone root-mean-square deviation (P-RMSD) metrics across all conformations. Minor groove widths were measured using the 3DNA software package^60^. The resulting outputs from 3DNA were processed using a Python script previously provided in our GitHub repository (https://github.com/CSB-KaracaLab/NucDNADynamics). For each simulation set, essential statistics including mean and standard deviation values for minor groove widths were computed.

For the calculation of P-RMSD values, idealized B-DNA reference structures corresponding to each studied sequence (CGC, CGG, CGT, and CGA motifs) were generated via the 3DNA web server. The P-RMSD was calculated specifically over the phosphorus atoms of the six nucleotides spanning the flipped cytosine region (5′-TGCGCT-3′). All simulation-derived DNA conformations were structurally aligned to their respective B-DNA reference structures and computed P-RMSD values using the software Profit (Martin, A.C.R., http://www.bioinf.org.uk/software/profit/).

### Comparative Dynamic Analysis Framework

To systematically quantify protein–DNA interactions across different sequence contexts and elucidate sequence-specific recognition mechanisms of DNMT3 enzymes, we developed the Comparative Dynamics Analysis (CDA) framework. CDA integrates base and shape readout analyses into a structured pipeline, enabling parallel evaluation of direct (base-specific) and indirect (shape-dependent) recognition features. CDA consists of two main analytical modules (base and shape readout), each implemented through progressively detailed steps explained below:

#### Base-readout module

The base-readout analysis quantifies direct recognition of DNA bases by DNMT3 enzymes through base-specific hydrogen bonding.

*(i) Residue-level flanking nucleotide – amino acid interactions:* For each MD simulation involving DNMT3A/B-CGC/G/T/A complexes, intermolecular interactions were systematically computed. Interaction data frames from independent trajectories of the same system were merged within all steps of IDF. Interactions involving SAM-DNA and DNMT3-methylated cytosine were removed from the dataframes. Additionally, base-specific interactions were identified by excluding backbone atoms of amino acids and sugar-phosphate backbone atoms of nucleotides. The remaining hydrogen bonds were quantified for each complex. To identify key interactions in the flanking region, interactions within positions +1 to +3 of the DNA were isolated. The number of base-specific hydrogen bonds for each interacting pair was calculated. Interactions occurring in less than 10% of the simulation time (4×400 ns, 3204 frames total, and less than 320 hydrogen bonds) excluded. Interaction intensities for each complex were calculated by dividing the total number of hydrogen bonds by the total number of frames (3204) and visualized using heatmaps.
*(ii) Atom-level flanking nucleotide – amino acid interactions:* To achieve atomistic resolution of the base-specific hydrogen bond formation identified in Step (i), the interacted atoms of each pair were identified. The total number of hydrogen bonds formed in each atom pair was calculated, and cumulative numbers were visualized using barplots. This detailed atom-level analysis provided insights into the specificity and dynamics of base-specific hydrogen bond interactions within the flanking nucleotide – amino acid interface.

#### Shape-readout module

The shape-readout analysis characterizes indirect recognition through DNA shape deformations and electrostatic interactions at the protein–DNA interface.

*(i) Quantification of DNA deformation:* DNA deformation relative to idealized B-DNA was quantified by calculating P-RMSD values across the simulation trajectories. P-RMSD served as a dynamic measure for DNA backbone flexibility and structural deformation. Probability distributions of P-RMSD values were visualized as boxplots for comparative analysis of deformation trends across different sequence contexts and enzymes.
*(ii) Electrostatic interaction energy correlation:* Local electrostatic interaction energies at the protein–DNA interface were computed across simulation frames. Correlations between P-RMSD values and electrostatic energies were analyzed via scatter plots, with Pearson correlation coefficients indicating the strength of coupling between DNA backbone deformation and charge complementarity. This step identified sequences exhibiting significant shape-dependent recognition.
*(iii) Minor groove width profiling and residue-level upstream flanking nucleotide – amino acid interactions:* The minor groove width was tracked as a positional feature along the upstream regions (positions –1 to –3 upstream of the CpG site). Profiles of minor groove width were plotted to visualize sequence-dependent shape alterations via standard deviation and mean values. Residue-level interactions, including base-specific hydrogen bonds and hydrophobic contacts at these upstream flanking positions, were quantified and visualized using interaction heatmaps. Interaction intensities were again computed by applying the same threshold criterion (at least 10% occurrence) to ensure robust quantification, and interaction intensities for each complex were calculated by dividing the total number of hydrogen bonds by the total number of frames (3204).

All CDA analyses were implemented using custom scripts developed in Python libraries. Structural analyses and visualizations were conducted using PyMOL. Complete CDA scripts, methodologies, and raw simulation data are openly available at: https://github.com/CSB-KaracaLab/DNMT3AB_specificity

