## Supporting Information for "So close yet so far apart: Distinct flanking sequence recognition by DNMT3A and DNMT3B"

**Supplementary Information Content**

**Supplementary Figure S1:** Intermolecular base-specific hydrogen bond analysis of DNMT3A cognate and non-cognate systems for all replica simulations.

**Supplementary Figure S2:** Intermolecular base-specific hydrogen bond analysis of DNMT3B cognate and non-cognate systems for all replica simulations.

**Supplementary Figure S3:** Hydrogen accepting and donating capabilities of nucleotides (from major groove) and important amino acids.

**Supplementary Figure S4.** Comparative analysis of DNA backbone deformation and minor groove widths in DNMT3A and DNMT3B complexes.

**Supplementary Figure S5:** Pairwise ionic bond profiles for DNMT3A and DNMT3B cognate and non-cognate complexes.

**Supplementary Figure S6:** RMSF analysis of DNMT3A complexes.

**Supplementary Figure S7:** RMSF analysis of DNMT3B complexes.

**Supplementary Figure S8:** Comparative structural flexibility analysis of DNMT3A and DNMT3B enzymes.

**Supplementary Figure S9:** RMSD analysis of DNMT3A and DNMT3B systems.

**Supplementary Movie 1:** Dynamic hydrogen bond formation that drives sequence selectivity of DNMT3A and its cognate CGC motif. The stable hydrogen bonds formed by Arg836 of DNMT3A with the G+2' are highlighted.

**Supplementary Movie 2:** Dynamic hydrogen bond formation that drives sequence selectivity of DNMT3B and its cognate CGG motif. Lys777 forms hydrogen bonds with G+1 and G+2, while Asn779 compensates for the loss of branching in Lys777 by forming hydrogen bonds with C+2'.

**Supplementary Figures**

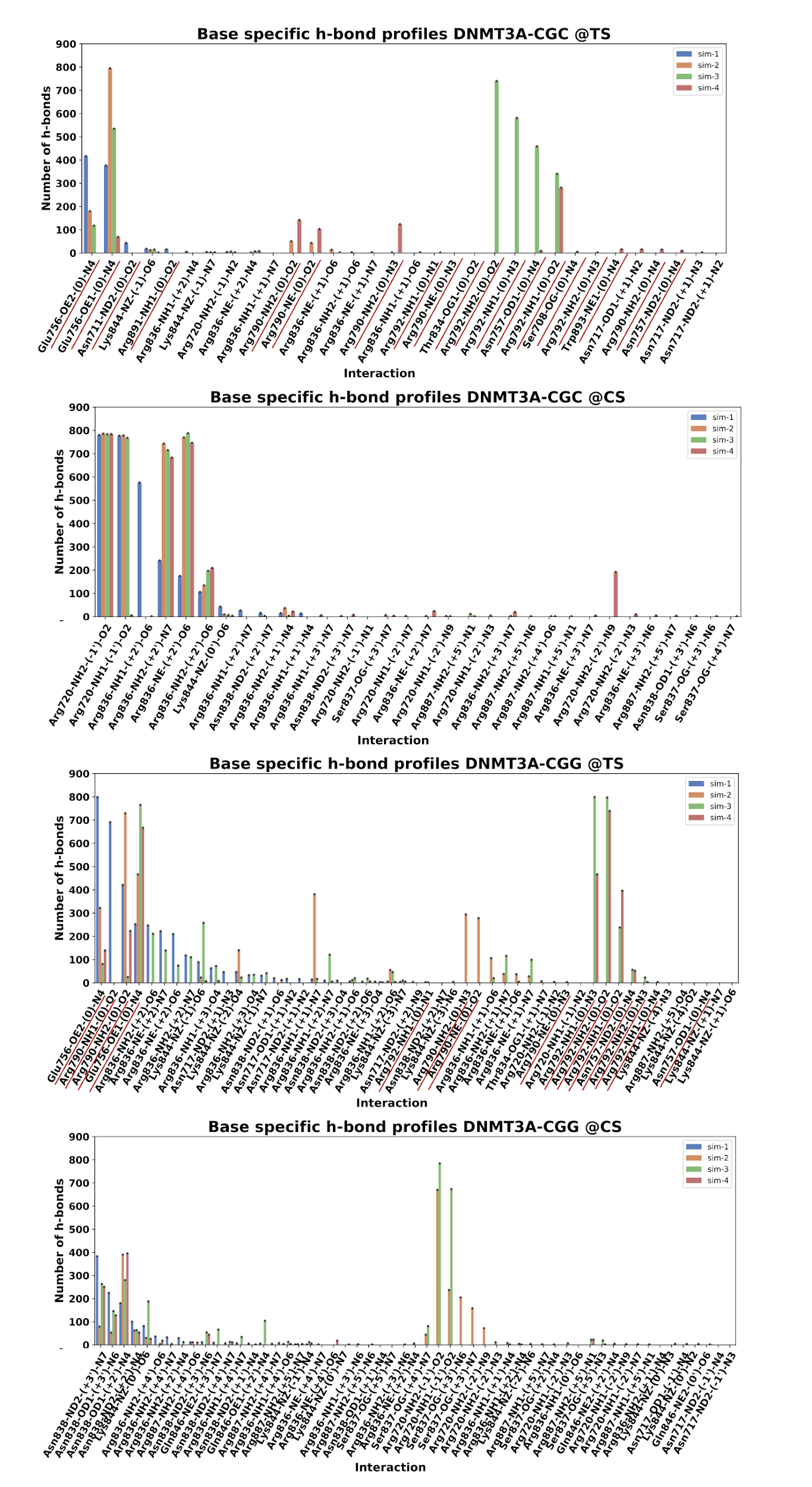

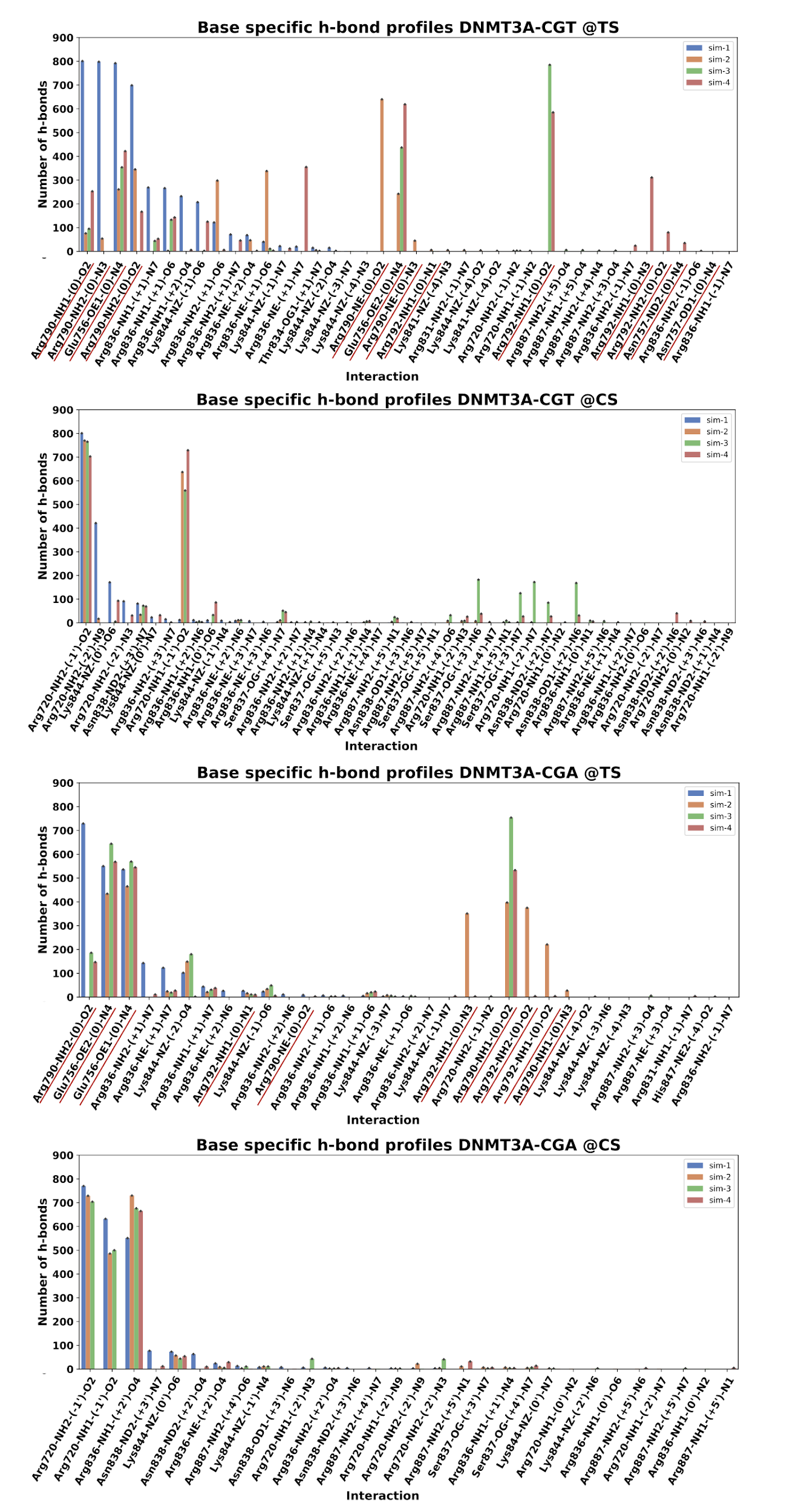

**Supplementary Figure S1: Intermolecular base-specific hydrogen bond analysis of DNMT3A cognate and non-cognate systems for all replica simulations.** Atomistic intermolecular base-specific hydrogen bond analysis is presented for each replica of molecular dynamics simulation involving DNMT3A cognate and non-cognate systems. In the plots, "TS" denotes the target strand of DNA, while "CS" represents the complementary strand interactions of the DNA. Each bar illustrates the interaction number observed during the production run (400 ns, N=801 conformations). The x-axis of the plot provides details about the interacting protein residue, residue side chain atom, nucleotide position, and nucleotide base atom. The interactions formed with the flipped cytosine are underscored with red.

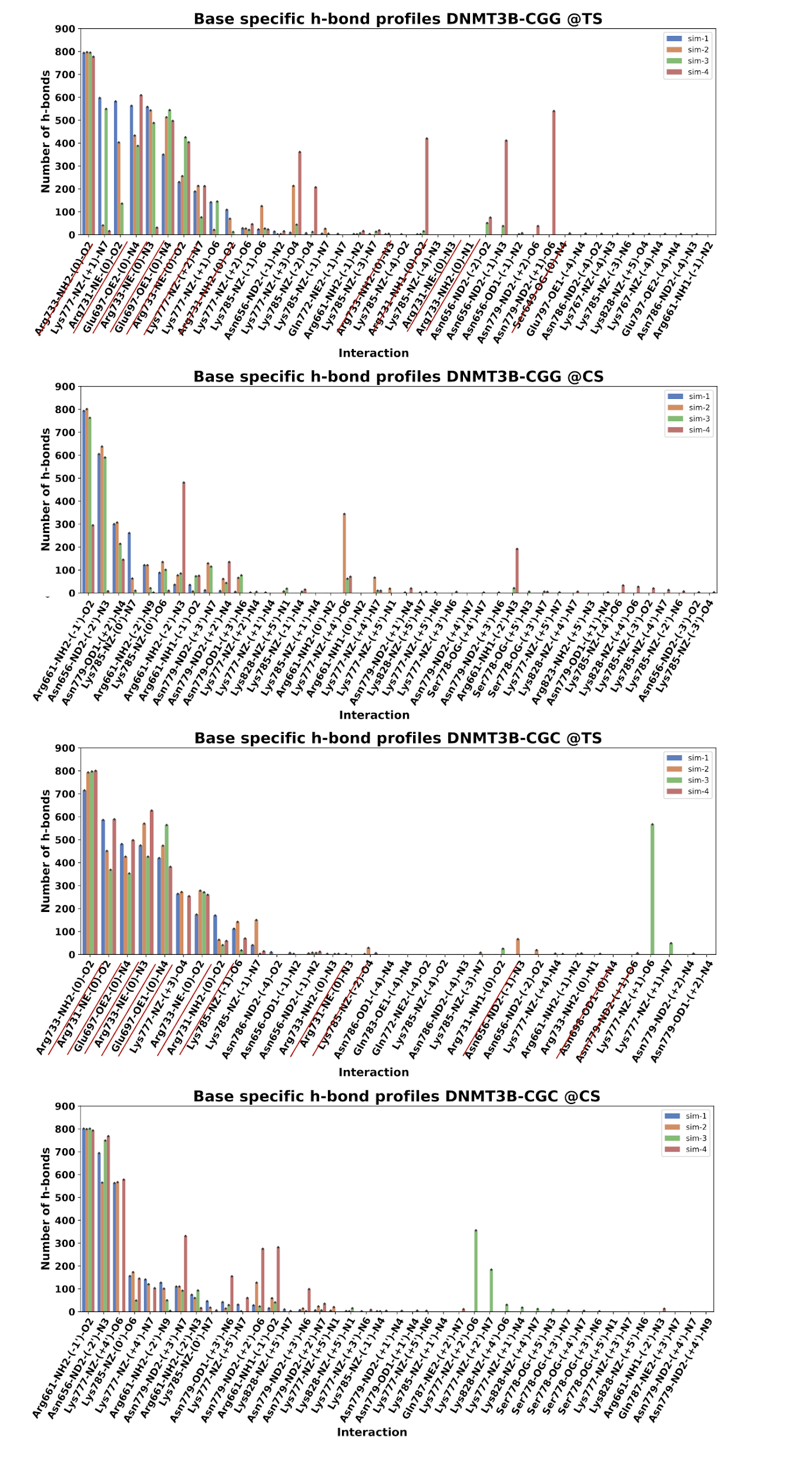

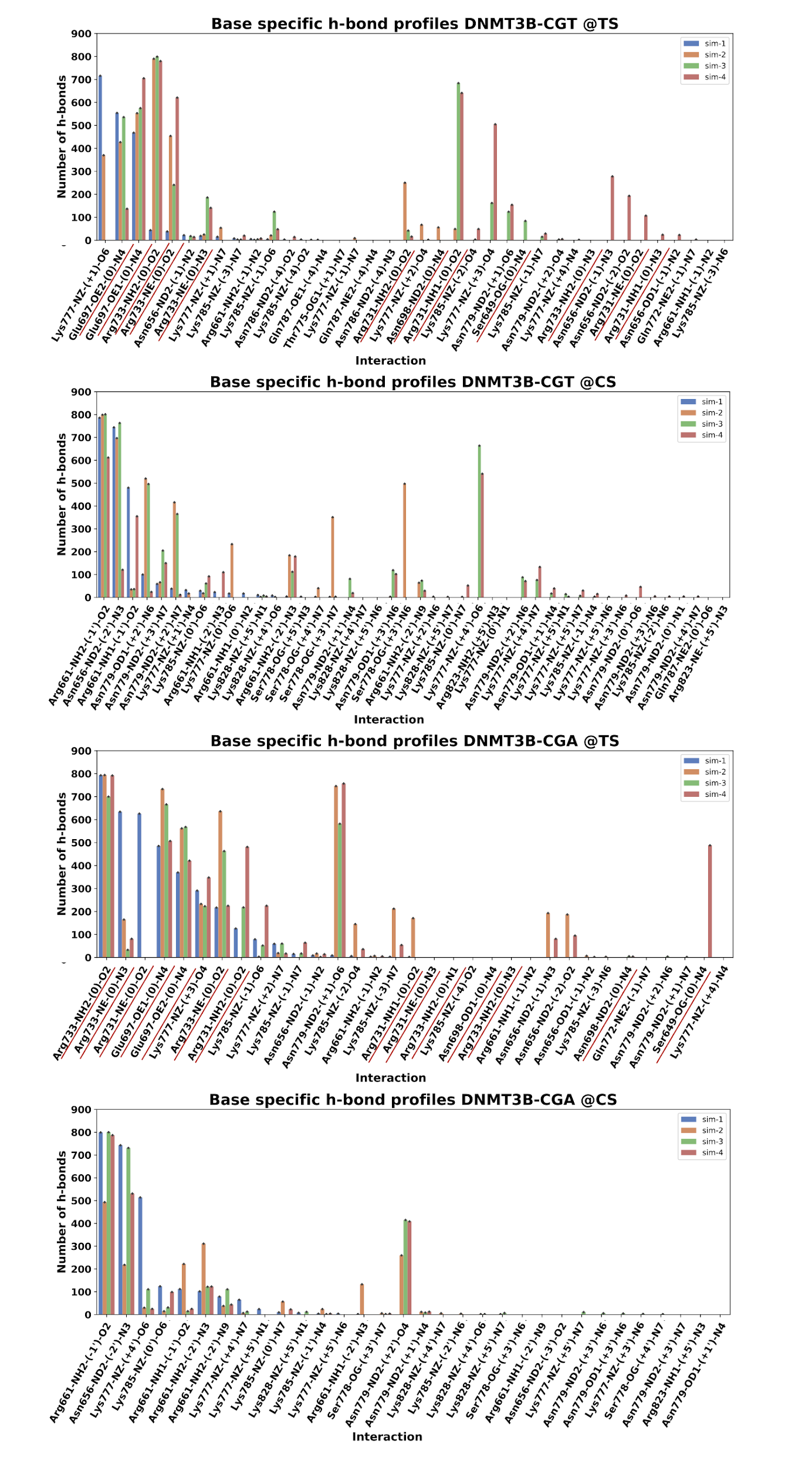

**Supplementary Figure S2: Intermolecular base-specific hydrogen bond analysis of DNMT3B cognate and non-cognate systems for all replica simulations.** Intermolecular base-specific hydrogen bonds at the atomistic scale are presented for each replica simulation involving DNMT3B cognate and non-cognate systems. In the plots, "TS" denotes the target strand of DNA, while "CS" represents the complementary strand interactions of the DNA. Each bar illustrates the interaction number observed during 400 ns production run (N=801 conformations). The x-axis of the plot provides details about the interacting protein residue, residue side chain atom, nucleotide position, and nucleotide base atom. The interactions formed with the flipped cytosine are underscored with red.

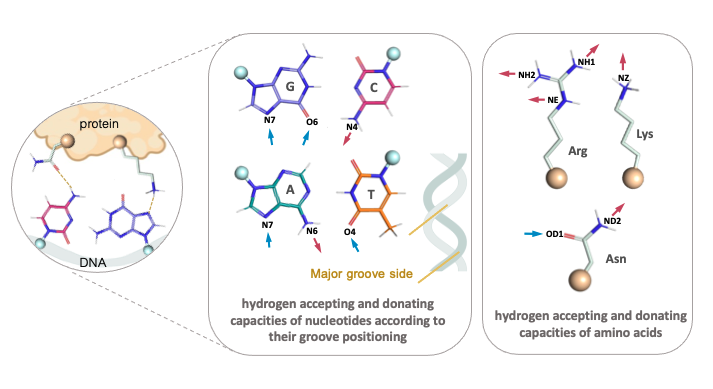

**Supplementary Figure S3: Hydrogen accepting and donating capabilities of nucleotides (from major groove) and important amino acids.** Blue arrows represent hydrogen-accepting, where red arrows show hydrogen-donating positions. Blue spheres represent the attachment point of the bases to the backbone; gold spheres represent the attachment point of side chains to the backbone. Through the major groove: Guanine accepts two hydrogens at N7 and O6, cytosine donates a hydrogen from N4, adenine accepts and donates hydrogens at N7 and from N6, thymine accepts a hydrogen at O4. Arg and Lys donate hydrogen from their NH1, NH2, NE, and NZ atoms, while Asn both accepts and donates hydrogens at its OD1 and from ND2 atoms, respectively.

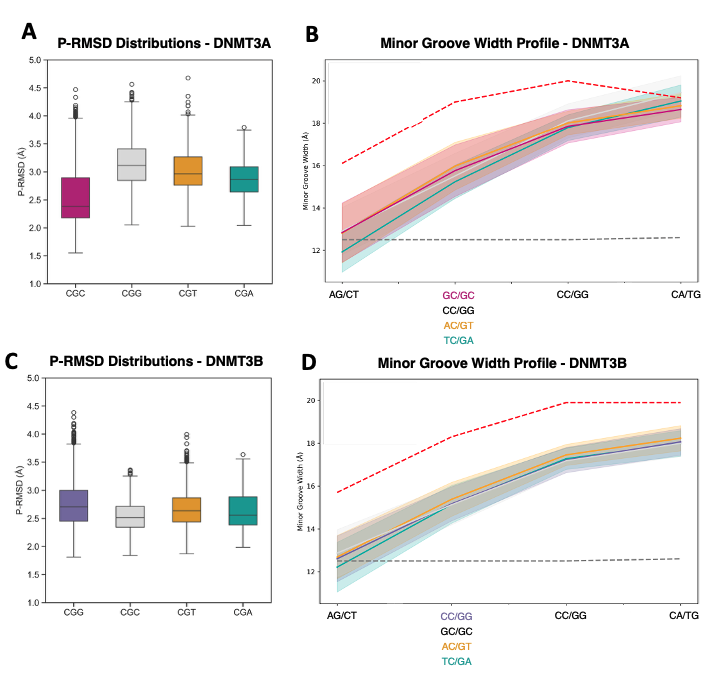

**Supplementary Figure S4. Comparative analysis of DNA backbone deformation and minor groove widths in DNMT3A and DNMT3B complexes. A)** P-RMSD distribution of
(**A**, **C**) Boxplots represent distributions of phosphate backbone RMSD (P-RMSD, Å) relative to ideal B-DNA across different flanking sequence contexts for (**A**) DNMT3A and (**C**) DNMT3B complexes, calculated over the MD simulation trajectories. Outliers are indicated by circles. (**B**, **D**) DNA minor groove width profiles (Å) averaged along the sequence flanking the CpG site in (**B**) DNMT3A (**D**) and DNMT3B complexes. Solid lines represent mean values, and shaded areas denote standard deviation across the trajectory. Red dashed and grey dashed lines indicate the minor groove of crystal structures and minor groove of B-DNA typically observed in DNA structures, respectively. Different colors correspond to specific sequence contexts as labeled in each panel.

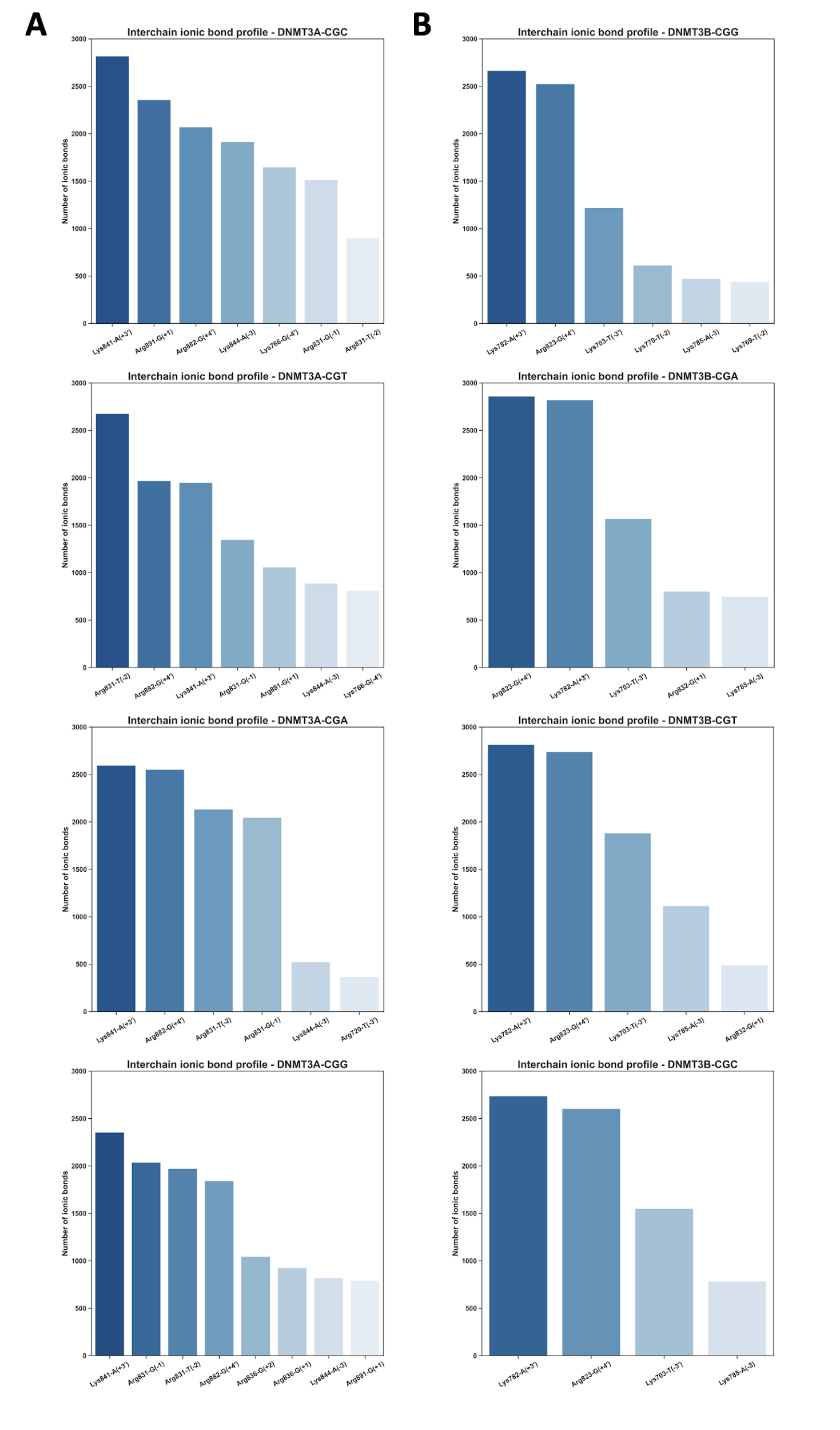

**Supplementary Figure S5: Pairwise ionic bond profiles for DNMT3A and DNMT3B cognate and non-cognate complexes.** Atom-based ionic bond analysis of **(A)** DNMT3A-CpG, and **(B)** DNMT3B-CpG motifs. Each bar shows the interaction number during 1.6 µs molecular dynamics simulations. The x-axis of the plot shows the interacting protein residue and nucleotide position.

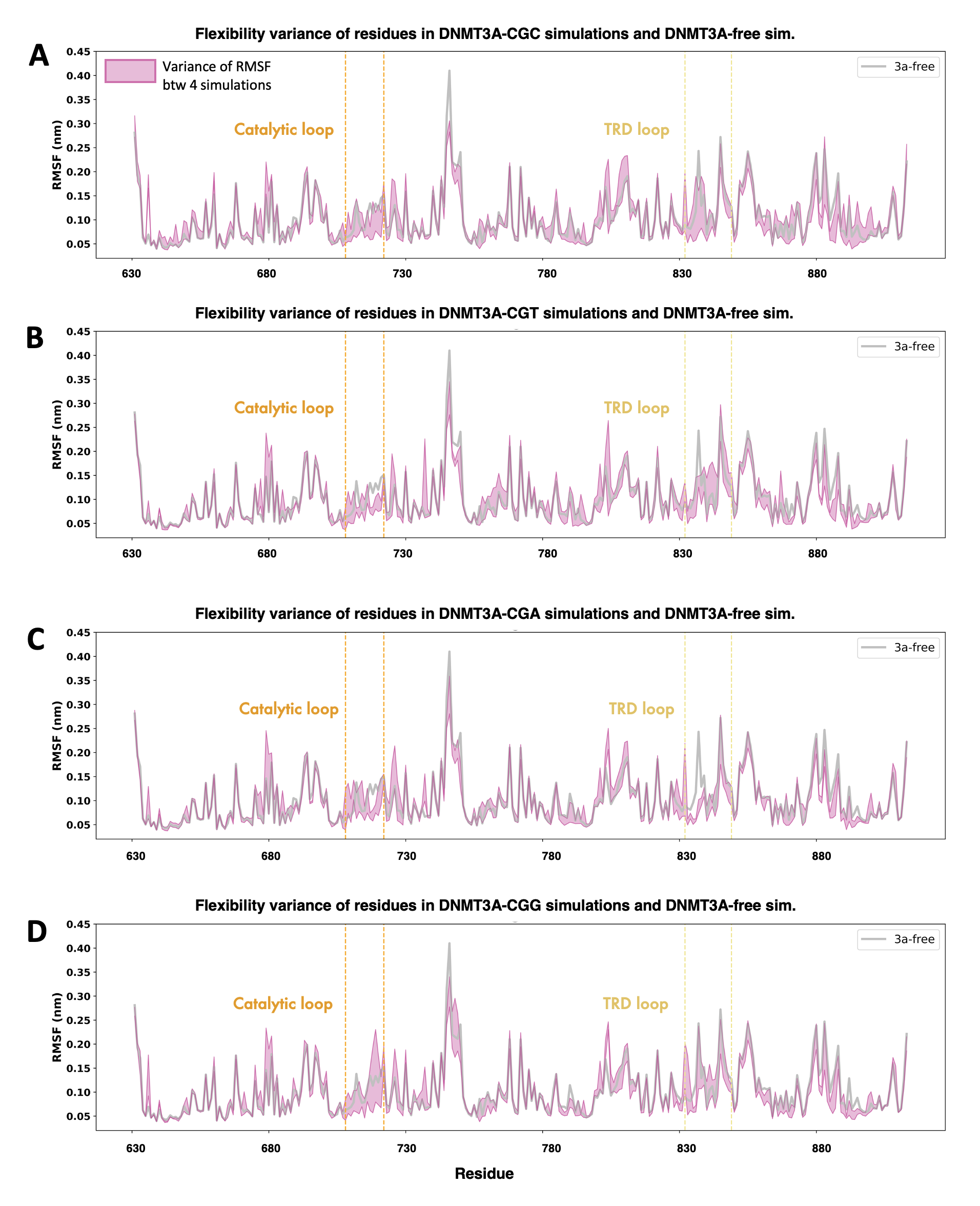

**Supplementary Figure S6: RMSF analysis of DNMT3A complexes.** Root Mean Square Fluctuation (RMSF) values of **(A)** DNMT3A-CGC, **(B)** DNMT3A-CGT, **(C)** DNMT3A-CGA, and **(D)** DNMT3A-CGG complexes calculated by subtracting 100 ns equilibrium time from 500 ns simulation. The orange lines represent the amino acid range of the catalytic loop, while the yellow lines represent the amino acid range of the TRD loop. The shaded areas depict the variance between simulations across four replicates. Gray lines show the RMSF values for the apo form of DNMT3A.

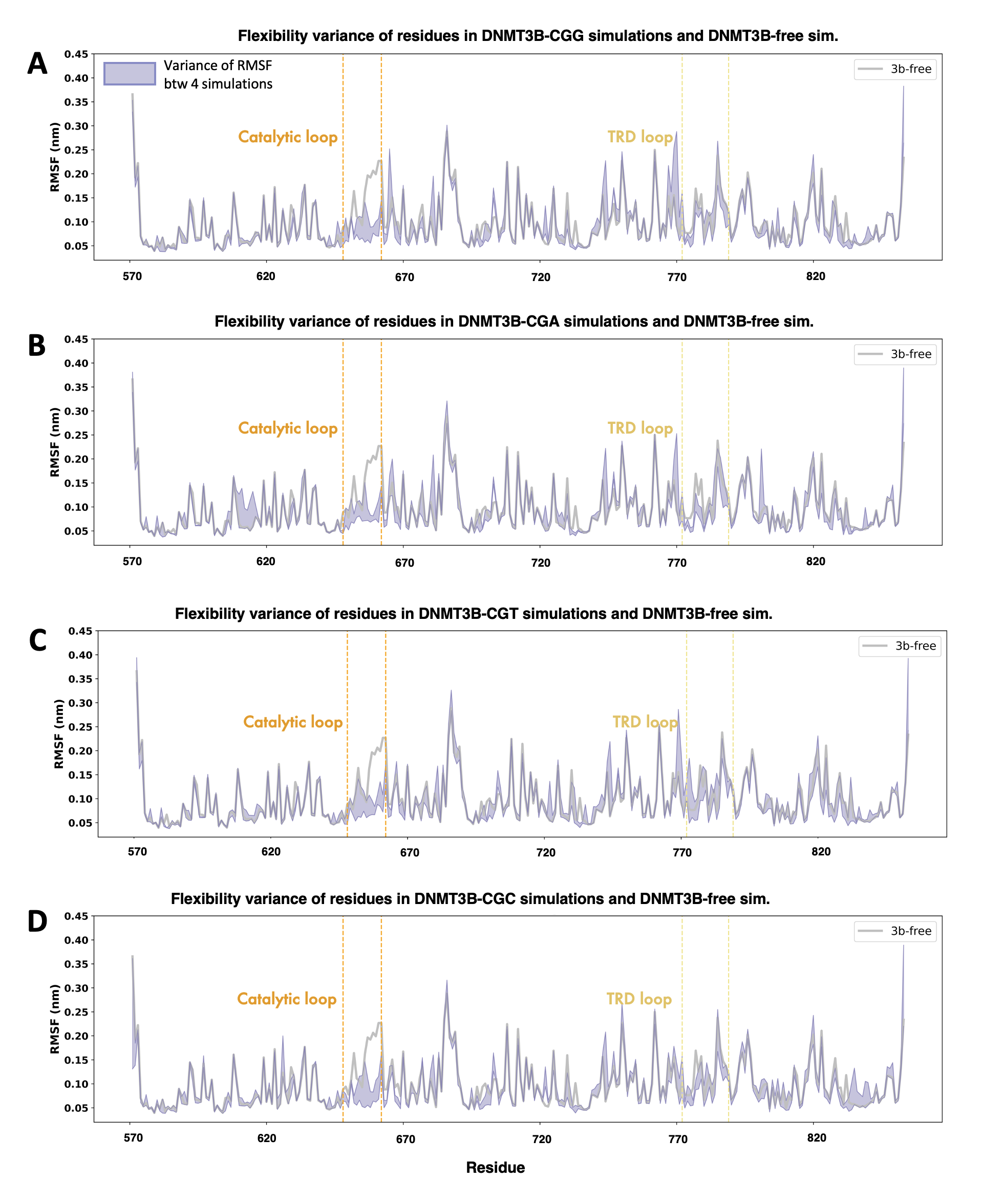

**Supplementary Figure S7: RMSF analysis of DNMT3B complexes.** Root Mean Square Fluctuation (RMSF) values of **(A)** DNMT3B-CGG, **(B)** DNMT3B-CGA, **(C)** DNMT3B-CGT, and **(D)** DNMT3B-CGC complexes calculated by subtracting 100 ns equilibrium time from 500 ns simulation. The orange lines represent the amino acid range of the catalytic loop, while the yellow lines represent the amino acid range of the TRD loop. The shaded areas depict the variance between simulations across four replicates. Gray lines show the RMSF values for the apo form of DNMT3B.

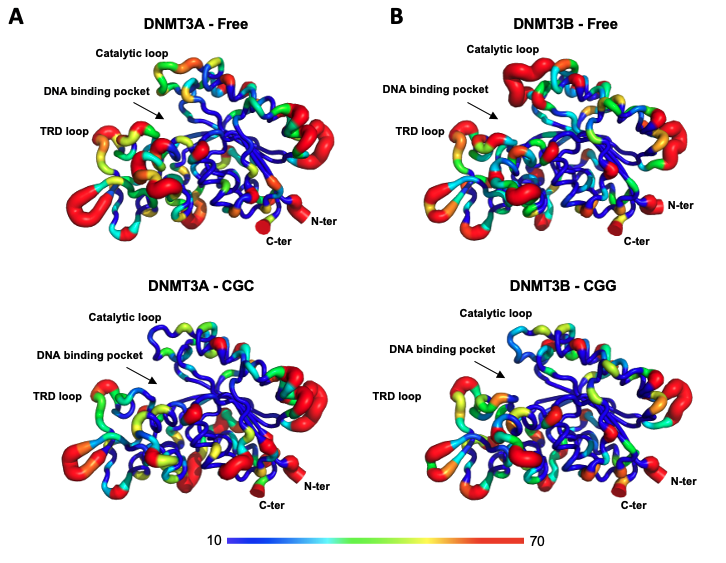

**Supplementary Figure S8: Comparative structural flexibility analysis of DNMT3A and DNMT3B enzymes.** Tube thickness and color represent residue-level flexibility (B-factor values, Å²) averaged over MD trajectories of DNMT3A (**A**) and DNMT3B (**B**) in free (unbound) and DNA-bound forms. Flexibility scale ranges from low (blue, rigid) to high (red, flexible). Key structural regions involved in DNA recognition and catalysis—including the target recognition domain (TRD) loop, catalytic loop, and DNA-binding pocket—are highlighted.

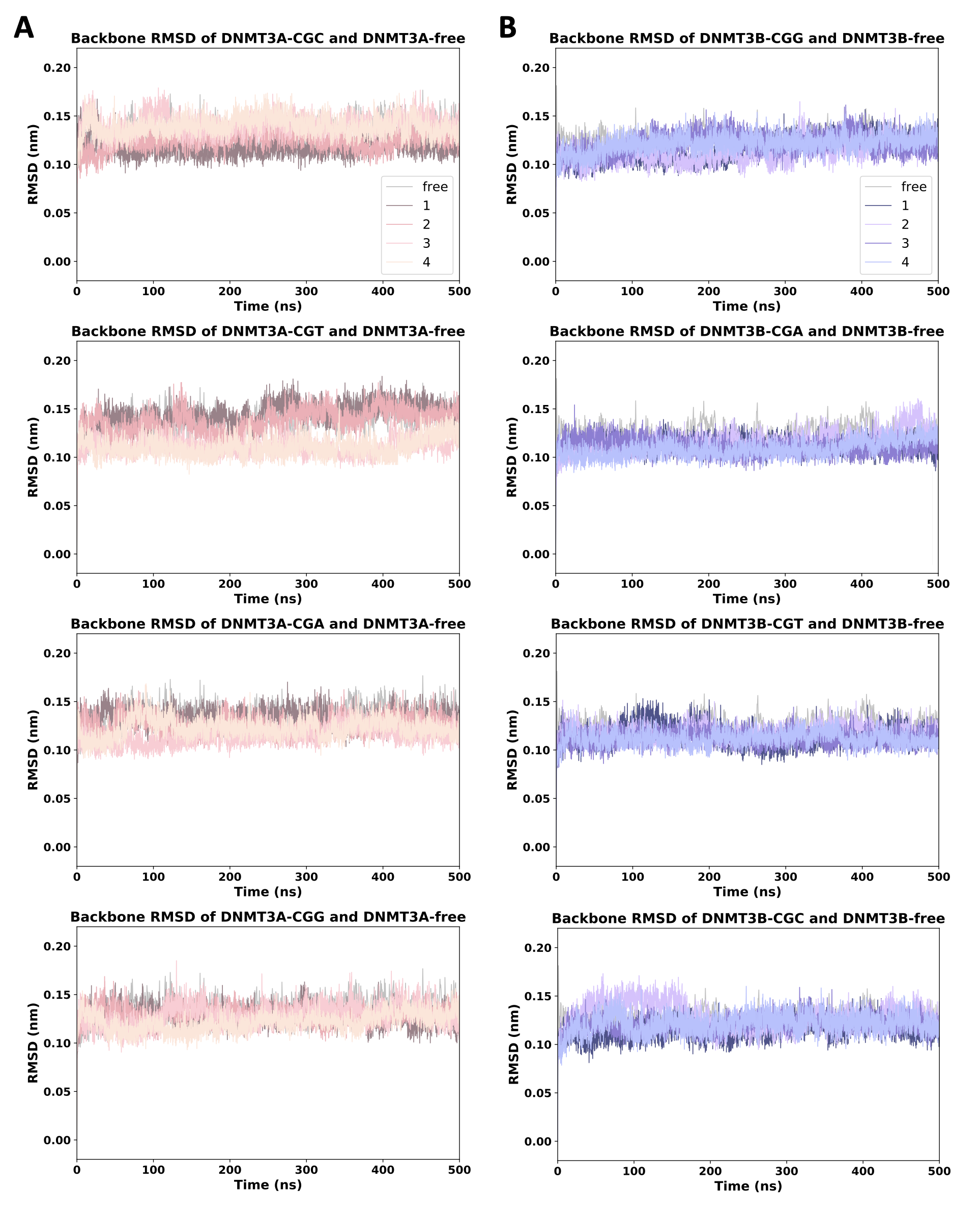

**Supplementary Figure S9: RMSD analysis of DNMT3A and DNMT3B systems.** Root Mean Square Deviation (RMSD) values calculated over backbone atoms of the systems during 500 ns molecular dynamics simulations according to their initial conformations. The RMSD values from four replicated simulations are presented for **(A)** DNMT3A-CGC/T/A/G and **(B)** DNMT3B-CGG/A/T/C complexes. The gray lines indicate the RMSD values for the apo forms of DNMT3A and DNMT3B enzymes.
